# Measuring spectrally-resolved information transfer for sender- and receiver-specific frequencies

**DOI:** 10.1101/2020.02.08.939744

**Authors:** Edoardo Pinzuti, Patricia Wollsdtadt, Aaron Gutknecht, Oliver Tüscher, Michael Wibral

## Abstract

Information transfer, measured by transfer entropy, is a key component of distributed computation. It is therefore important to understand the pattern of information transfer in order to unravel the distributed computational algorithms of a system. Since in many natural systems distributed computation is thought to rely on rhythmic processes a frequency resolved measure of information transfer is highly desirable. Here, we present a novel algorithm, and its efficient implementation, to identify separately frequencies sending and receiving information in a network. Our approach relies on the invertible maximum overlap discrete wavelet transform (MODWT) for the creation of surrogate data in the computation of transfer entropy and entirely avoids filtering of the original signals. The approach thereby avoids well-known problems due to phase shifts or the ineffectiveness of filtering in the information theoretic setting. We also show that measuring frequency-resolved information transfer is a partial information decomposition problem that cannot be fully resolved to date and discuss the implications of this issue. Last, we evaluate the performance of our algorithm on simulated data and apply it to human magnetoencephalography (MEG) recordings and to local field potential recordings in the ferret. In human MEG we demonstrate top-down information flow in temporal cortex from very high frequencies (above 100Hz) to both similarly high frequencies and to frequencies around 20Hz, i.e. a complex spectral configuration of cortical information transmission that has not been described before. In the ferret we show that the prefrontal cortex sends information at low frequencies (4-8 Hz) to early visual cortex (V1), while V1 receives the information at high frequencies (> 125 Hz).

**Author Summary:** Systems in nature that perform computations typically consist of a large number of relatively simple but interacting parts. In human brains, for example, billions of neurons work together to enable our cognitive abilities. This well-orchestrated teamwork requires information to be exchanged very frequently. In many cases this exchange happens rhythmically and, therefore, it seems beneficial for our understanding of physical systems if we could link the information exchange to specific rhythms. We here present a method to determine which rhythms send, and which rhythms receive information. Since many rhythms can interact at both sender and receiver side, we show that the interpretation of results always needs to consider that the above problem is tightly linked to partial information decomposition - an intriguing problem from information theory only solved recently, and only partly. We applied our novel method to information transfer in the human inferior temporal cortex, a brain region relevant for object perception, and unexpectedly found information transfer originating at very high frequencies at 100Hz and then forking to be received at both similarly high but also much lower frequencies around 20Hz. These results overturn the current standard assumption that low frequencies send information to high frequencies.

## Introduction

Many natural or artificial complex systems perform distributed computation. In a distributed computation multiple relatively simple parts of the system perform rather elementary operations on their inputs, but do communicate heavily amongst each other in order to jointly implement complex computations. Thus, to understand these joint computations, measuring the information transferred between the parts of the system is crucial. A mathematically rigorous measure of information transfer is the transfer entropy (TE) [1]. TE, as a model-free information theoretic measure, is ignorant of the details on how the information transfer is physically implemented, which is indeed a highly desirable property when we only want to detect and measure information transfer. However, many systems display highly rhythmic activity when performing distributed computation, suggesting that measuring the information transfer associated with different spectral components may provide valuable additional insights. This holds in particular for biological neural systems where rhythmic or quasi-periodic activity is found frequently across many scales from spiking activity of individual neurons to electroencephalographic (EEG) recordings of large pools of neurons (see [2] and references therein).

Early attempts [3] to obtain the desired frequency-resolved measurement of TE resorted to narrow-band filtering of the data from information source and information receiving target and to feeding the resulting narrow-band signals into a TE analysis. Yet, these approaches come with certain problems that are well-known from the field of Granger-Causality (GC) analysis. Due to the equivalence of GC and TE for jointly Gaussian variables [4], these problems carry over to TE analyses.:

1. Most importantly, the use of filters prior to TE computation for achieving frequency resolution will lead to false positive results due to phase distortions, or will not have the desired frequency-specific effect at all, i.e. TE computed from filtered and unfiltered signals is approximately the same. This latter effect is due to the fact that reducing the power of a signal does not reduce the information contained in it, except for additional effects of signal quantization. Both modes of failure are well known from results on the linear approximations of TE (e.g. via Granger causality, [4–7]).
2. The usual focus on information transfer between a source and a target within a specific narrow frequency band (driven by ideas of synchronization) practically confines the analysis to the linear interaction regime—even when using a nonlinear, model-free, measure like TE. This is because many interesting nonlinear mechanisms of information transfer will actually transform frequencies between source and target.
3. Within-frequency band analyses also ignore the potential many-to-many relationships that source and sender frequencies could have when there is information transfer. For example, a signal at approximately 10 Hz in the source may not seem to transfer information to any specific frequency of the target, yet when considering all frequencies of the raw target signal together, then non-zero TE from 10 Hz at the source to the full signal at the target is observed. In the same way, source signals in two or more bands may have to be considered jointly to reveal TE to the target. On the other hand, multiple bands in the source may carry and transfer identical information to the target, that might be ‘double counted’ in a naive frequency-resolved analysis. Last, when one frequency is observed sending information and another receiving information, it is not guaranteed that this is actually the same information. In other words, information may be sent from the observed source frequency to all target frequencies jointly while *different* information may be sent from all source frequencies jointly to a specific target frequency.

To circumvent filtering-related problem 1 we here suggest a novel algorithm to obtain frequency resolution of TE without ever filtering the original signals. Instead of filtering the original signals, we apply filtering in the creation of surrogate data representing the null-hypothesis of no information transfer at the frequencies of interest. This way, we use the potential distorting effect of filtering to our advantage, and destroy temporal order in the surrogates instead of changing the spectra of the signals. To solve problem 2, we create the frequency-specific surrogate data separately for source- and target-frequencies. This reflects that frequency specific TE is a many-to-many problem, and that within frequency-band analyses may miss most or all of the information transfer. We then discuss problem 3 at the conceptual level, and we explain how splitting of source and target signals into multiple frequencies means that one has to deal with a multivariate problem that is of the ‘partial information decomposition’ (PID)-type. As the PID problem is of considerable complexity and keeps challenging mathematicians to date, we cannot present a full solution, but rather use the PID formalism to further elucidate our approach to frequency-resolve source and target separately. This restrained approach should serve as a reminder of the often overlooked complexity of the above problem.

## Theory, Algorithm and Methods

We start this section with some technical background material on the transfer entropy measure and on the creation of frequency-specific surrogate data in which only a single spectral component has been altered. These two technical sections maybe be skipped at first reading or when not interested in technical details. After this technical background we then present the two core algorithms of this study, which serve to identify source-frequency specific and receiver-frequency specific information transfer.

### Background

#### Technical Background: Transfer Entropy and Multivariate Transfer Entropy

Transfer entropy (TE) as the fundamental measure of information transfer was introduced first in [1] in a bivariate framework for two random processes 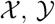 as:

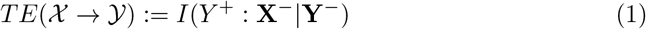

where *I*(·: ·|·) is the conditional mutual information and *Y* +, Y^-^, X^-^ are, respectively, a future random variable of the process 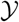, a vector of suitably chosen past random variables of the past of that process, and a suitably chosen vector of past random variables of the process 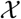 (see [1,8–11] for considerations on the correct choice of the past random variables). TE measures the amount of information transferred between a single source and a single target process. In a network setting where multiple sources may interact to transfer information to a target, or where multiple sources transfer information redundantly to a target, TE needs to be extended to a multivariate formalism in order to avoid spurious results. Thus, we need to measure the information transfer from a single source to a target, but now in the context of all other relevant sources in an observed network [12,13]. In other words, the multivariate TE (mTE) for a system of *M* random processes 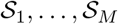, observed over *D* discrete time steps, represented by random vectors **S**_*i*_ = (*S*_*i*,1_, …, *S*_*i,D*_), measures the information transfer as a *conditional* mutual information of the following form:

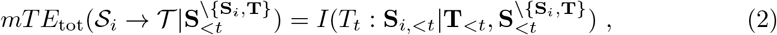

where 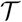 is a process considered as the current target of the information transfer, **S**_*i*,<*t*_ = (*S_i,t–δ_*,…, *S_i,t–k_*), with *δ* ≤ *k* is a vector of random variables chosen from **S**_*i*_ from the past of the current time point *t*. The last element *k* of this vector is chosen such that **S**_*i,<t*_ renders *T_t_* conditionally independent of all variables in the process **S**_*i*_ that are further back in time. The delay parameter *δ* is chosen such that it reflects the physical delay in the system (see [10]). 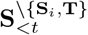 signifies the collection of past random variables from all other processes except **S**_*i*_ and *T*. Last, the subscript ‘tot’ means that we are focusing on the total information transferred from all relevant past variables of the process **S**_*i*_, rather than the contribution of each individual variable **S**_*i,d*_.

Computing the *mTE_tot_* in equation 2 exactly is an NP-hard problem [14], and approximations are necessary for practical use. This problem was recently addressed in [15,16], with the implementation of an approximate greedy algorithm in the IDT^*xl*^ toolbox, which allows a large-scale directed network inference with mTE [11] and is freely available from GitHub (https://github.com/pwollstadt/IDTxl). IDTxl performs a greedy algorithm with an iterative sequence of statistical steps to infer the ‘relevant’ sources of the network, thus reducing the dimensionality of the problem, and allows to properly construct the nonuniform embedding of source and target time-series [17,18], i. e. it also yields approximations for parameters like *δ* and *k*.

The *mTE_tot_* estimated from the original data will be the test statistic of interest for our spectral mTE algorithm in which it is statistically tested against a null distribution of the same measure computed from frequency-specific surrogate data constructed by the maximum overlap discrete wavelet transform (MODWT).

In the exposition of the spectral mTE algorithm below we will assume that the *mTE_tot_* estimation for the original data has been performed and a set of sources **S**_*i,<t*_ significantly contributing *mTE* for the target 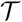 has been identified. This set of significant source processes with respect to a target will be passed to the spectral TE algorithm, to identify TE relations at specific frequencies in these sources and the target. The algorithm is applied to find the spectral components of contributing to the overall information transfer one source at a time **S**_*i,<t*_, but is of course repeated over all relevant sources of a target.

#### Technical Background: Maximum Overlap Discrete Wavelet Transform

Several methods have been established for surrogate data creation, each with its own limitations and advantages (see [19] for a review). Among many, wavelet-based methods allow to create frequency-specific surrogate data through randomization of the wavelet coefficients [20]. In particular, wavelet-based surrogates that preserve the local mean and the variance of the data were introduced by [21]. Similarly to [22], we employ the Maximal Overlap Discrete Wavelet Transform (MODWT), to transform the data in the wavelet domain. The MODWT is well defined for time-series of any sample size and produces wavelet coefficients and spectra unaffected by the transformation. [22].

The MODWT of a time-series *X* = (*X*_0_,…, *X*_*N*–1_) of *J*_0_ levels, where *J*_0_ is a positive integer, consists of *J*_0_ + 1 vectors: *J*_0_ vectors of wavelet coefficients 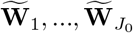 and an additional vector 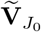 of scaling coefficients, all with dimension *N* (our exposition of the MODWT closely follows that of [23], pages 159-205). The coefficients of 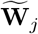 and 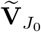 are obtained by filtering X, namely:

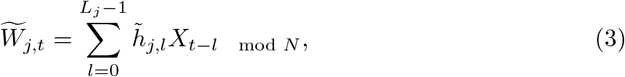

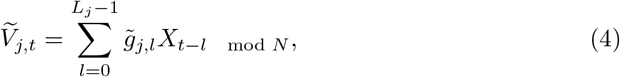

where 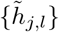 and 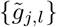 are the *j*th level MODWT wavelet and scaling filter, with *l* = 1,…, *L* being the length on the filter and *L_j_* = (2^*j*^ – 1)(*L* – 1) + 1. We can write the above in matrix notation as:

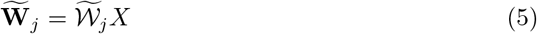

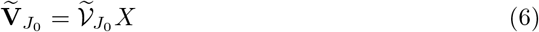

where each row of the *N* × *N* matrix of 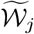 has values denoted by 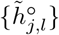, while 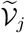 has values denoted by 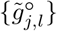, where 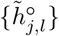 and 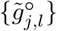 are the periodization of 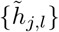 and 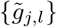 to circular filter of length *N* [23]. Thus, the MODWT treats *X* as if it were periodic, such periodic extension is known as ‘circular boundary condition’ [23]. Finally, the time series *X* can be retrieved from its MODWT with [23]:

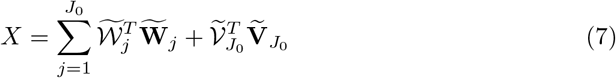

While, the coefficients 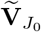 represent the unresolved scale [22,23], and capture the long term dynamics of *X*, the coefficients 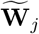 are associated with changes of the underlying dynamics, at a certain scale, over time. If *N* = 2^*j*^ and we set *J*_0_ = *J*, then a full decomposition is performed and the scale 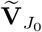 retains only the average constant of the data with all other information represented in the wavelet coefficients [22, 24]. Since in many applications a full decomposition is not necessary (e.g. the dynamic of a physical system is meaningful over a certain frequency range only), *J*_0_ can be set to any integer *J* ≤ ⌊(log_2_(*N*)⌋ so that the decomposition at any scale is shorter than the total length of the time series [25]. The selection of *J*_0_ determines the number of scales of resolution with the MODWT coefficients at a certain scale *j* related to the nominal frequency band |*f*| ∈ (1/2^*j*+1^,1/2^*j*^) [23]. Moreover, given 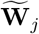 and 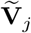 it is possible to reconstruct the time-series *X* through the inverse MODWT (IMODWT). If the coefficients are not modified, the IMODWT returns the original time-series *X* [23].

### Algorithms

#### Algorithm I: Identifying source- or receiver-frequency specific TE

##### Core idea

The core idea of the proposed algorithm is to never apply any frequency-specific signal processing to the original data from which TE is computed, as this is known to come with a whole host of problems [5, 7]. Rather, frequency-specificity is obtained by destroying TE-relevant signal properties (like temporal order) in a frequency-specific manner in *the surrogate data* and to then look for a significant drop in mTE in these surrogate data compared to the original mTE via non-parametric statistical testing. To this end, we create surrogate data via an invertible wavelet transform (maximum overlap discrete wavelet transform, MODWT) and a frequency (scale-) specific scrambling of the wavelet coefficients in time. Thus, in the surrogate data temporal order and phase relations are destroyed specifically in the band of interest, while the power spectra of the signals are preserved. As frequency separation is never perfect, we confine ourselves to only interpreting the wavelet-scale or frequency where the difference between the median of the distribution of mTE from surrogate signals and the value of total original multivariate TE (*mTE_tot_*, see next) is largest.

##### Implementation for source-frequency specific information transfer

As introduced above, we obtain a measure of frequency-specific information transfer by creating surrogate datasets in which the temporal ordering of the signals has been destroyed for specific spectral components of these signals—by first transforming into the frequency domain, then scrambling wavelet coefficients for a specific frequency and last transforming back to the time domain to obtain a surrogate dataset. Naively one may be tempted to apply this process to source and target processes at the same time. Yet, this approach would limit the analysis to within-band effects. As laid out in the introduction and also detailed in section *Frequency resolved TE as a partial information decomposition problem,* this would ignore the multivariate nature of the problem. Therefore, we apply the creation of frequency-specific surrogate data separately to source and target processes, i.e. we apply two variants of the analysis—one measuring source-frequency specific information transfer and the other measuring target-frequency specific information transfer. A combination of the results of both analyses is sometimes possible when carefully considering before the multivariate nature of the problem and prior knowledge (see section *Relation of the partial-information decomposition framework and the SOSO-algorithm* below).

We will now detail the algorithm variant for the measurement of source-frequency specific information transfer and report the relevant differences for measuring target-frequency specific information transfer afterwards.

Measuring source-frequency specific mTE relies on five main steps:

1. Perform a wavelet decomposition of the source time series through the MODWT to obtain a time-frequency representation of **S**_*i*_ in *J*_0_ scales.
2. At the j^th^ scale of the MODWT decomposition shuffle the wavelet coefficients to destroy information carried by the scale (frequency band)
3. Apply the inverse wavelet transform, IMODWT, to get back the time representation of the time series
4. Compute the *mTE*’ between the surrogate source and the target, conditional on all other significant sources in the network.

a. Repeat from step 2 to 4 for a number of permutations to build a surrogate data distribution.
b. Repeat from step 1 to 4 for all *J*_0_ scales.
5. Test whether the original *mTE_tot_* is above the 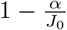 quantile of the surrogate-based distribution of *mTE’* values at each scale, i.e. perform a significance test with respect to the surrogate-derived distribution.

The operations implemented in the five steps are illustrated in Figure 1 and described in detail hereafter.

**Fig 1.**
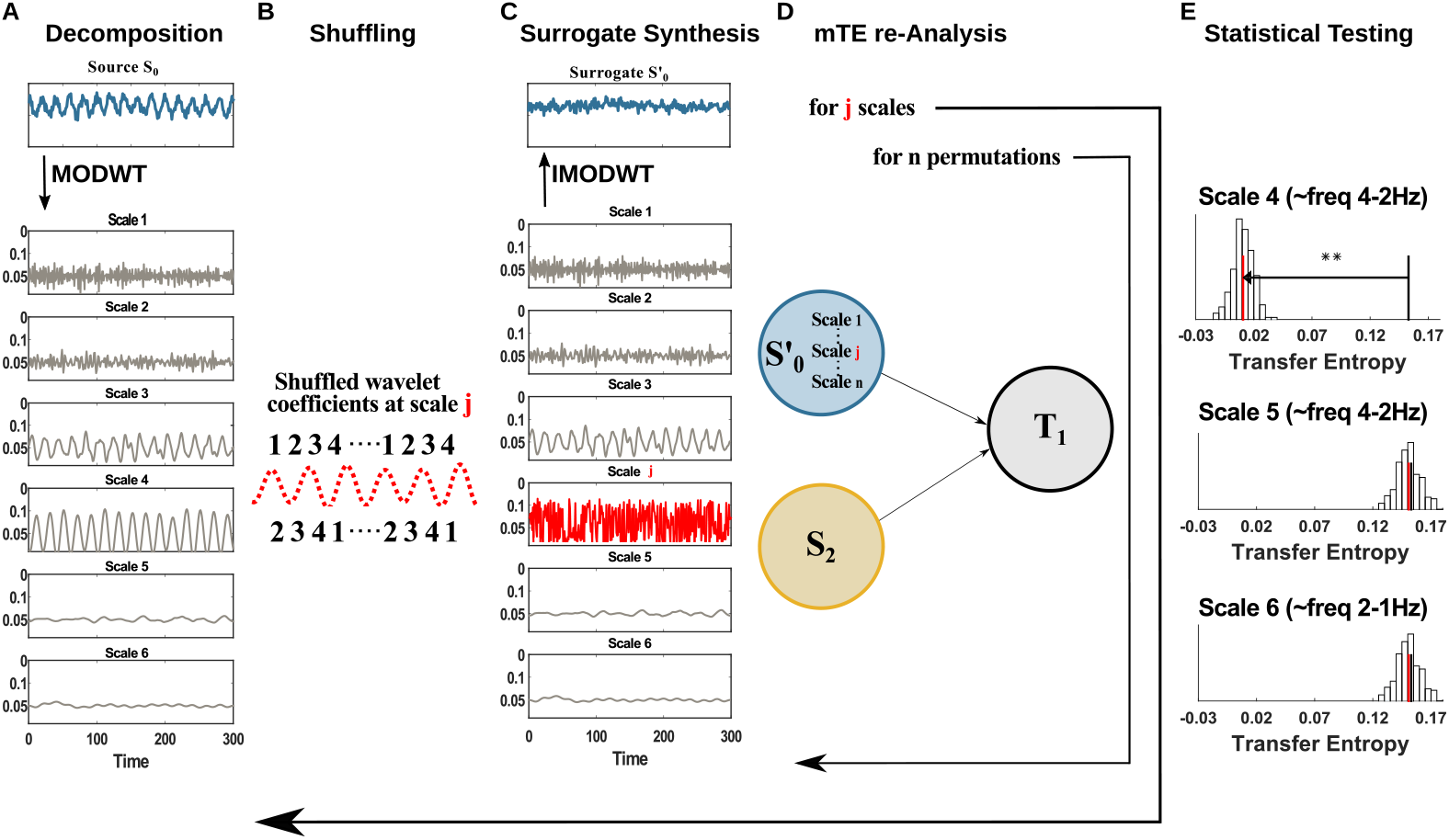
Spectral TE algorithm pipeline. (A) The neural signal (blue) is converted to a time-frequency representation (grey) using the invertible maximum overlap discrete wavelet transform (MODWT). (B) At a frequency (wavelet scale) of interest in the source (or the target) the wavelet coefficients are shuffled in time, destroying its connection to the target (or source). (C) The signal is recreated by the inverse MODWT. (D) The transfer entropy for the original and many shuffled signals is computed. (E) A statistical tests determines whether the shuffling reduced the information transfer, indicating that the transferred information was indeed encoded at the specific frequency. Each panel here shows the distribution of *mTE’* values (vertical bars) obtained from surrogate data where the wavelet coefficients of the scale of interest were shuffled, the median of this distribution (red line), and the original transfer entropy (black line). The analysis and the testing is repeated for all scales of interest (here 4,5,6).

**Step 1:** The source time-series is decomposed once into *J*_0_ scales through the MODWT (Fig. 1, Panel A). As introduced in section *Maximum Overlap Discrete Wavelet Transform* this decomposition gives a set of details coefficients 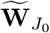 and an additional set of approximation coefficients 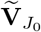. The latter is saved in this first step and utilized only in step 3, without any modification. Only the 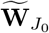 coefficients at the *j*^th^ scale under analysis are subjected to step 2. The current implementation uses a Least Asymmetric Wavelet (LA) as mother wavelet of length 8 or 16, since both lengths showed to be robust against spectral leakage and do not relevantly suffer from boundary-coefficient limitations. [21,23,26]. The creation of surrogate data for subsequent statistical testing comprises of the following steps 2 and 3.

**Step 2:** The frequency-specific information transfer between source and target is destroyed by shuffling the 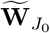 wavelet coefficients one scale at a time. The *j*^th^ scale under analysis is shuffled by randomly permuting the coefficients 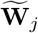, whereas all the other scales decomposed by the MODWT stay intact (Fig. 1, Panel B, *j*^th^ scale in red). We implement two alternative methods for the creation of surrogate data: a Block permutation of the wavelet coefficients [20] and the Iterative Amplitude Adjustment Fourier Transform (IAAFT) [20, 22]. Since there is no unique method of surrogate data creation and in many cases the employment of one method or another much depends on the specific analysis carried out by the user, we describe the two methods and the input parameters in section *Resampling Methods and the free parameters.*

**Step 3:** The unchanged set of coefficients, 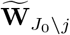, the unchanged 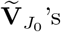, and the permuted coefficients at scale *j* 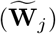 are submitted to the IMODWT, to reconstruct the surrogate source signal, 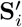, in the time-domain (Fig. 1, Panel C). This step is identical for both of the implemented surrogate-data creation methods: Block permutation of the wavelet coefficients and IAAFT. The reconstructed source 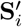 (*source surrogate*) differs from the source **S**_*i*_ only on the shuffled *j^th^* scale. In this way, we destroy the source-target information transfer only if the information transfer is carried by the j^th^ scale, otherwise the information transfer stays the same.

**Step 4:** With 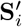 we compute again the *mTE_tot_* on the network previously identified. We illustrated this step in Fig. 1, Panel D. Let **S**_*i,<t*_ be the set of past variables of the *selected sources* and **T**_<*t*_, the past variables of the *selected target* previously found in the network analysis, with 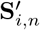 being the *n*-th *source surrogate* under analysis in the network at scale *j*; then, the *mTE*’ for the surrogate data is:

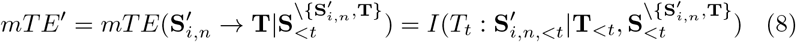

The algorithm is repeated from step 2 to step 4 for n *permutations*, with *n* = 1,…,*N*, to create a distribution of surrogate 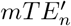 values; *N* is set according to the desired critical level for statistical significance (including Bonferroni correction for the number of scales, see below). Subsequently, all the *J*_0_ scales decomposed by the MODWT in step 1 are subjected to step 2, step 3 and step 4, such that *J*_0_ separate distributions of 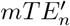-values, one for each scale, are obtained.

**Step 5:** As a final step, the *mTE_tot_* is tested for statistical significance against the *J*_0_ different distributions of *mTE’* surrogate values. If the 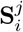 (where *j* is one of the scales decomposed by the MODWT) carries any information transfer to the target *T*, a significant drop of the *mTE*^’^ surrogates will be observed. This step is applied for all *J*_0_ scales under analysis and a Bonferroni correction is applied such that each individual scale is tested at the significance level *α/J*_0_.

Additionally, each scale analyzed is plotted, see Fig. 1, Panel E, and we restrict ourselves to interpret only the scale that shows maximal distance from the original *mTE_tot_*, 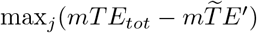, where 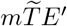 denotes the median of the surrogates distribution. We consider the maximal distance in addition to the statistical significance test because frequency decomposition is never perfect (e.g. leakage, noise and wavelet bands overlap). Indeed, validation of the algorithm on synthetic data (section 3) shows that the maximal distance reliably reflects the ground truth in the sender-receiver frequency information transfer, independently of the method employed for surrogate construction, whereas the statistical significance test can suffer from leakage effects on adjacent scales. Obviously, this limits the detectability of frequency-specific *mTE_tot_* to one source frequency and may be overly conservative. Thus, in scenarios, where information transfer from multiple sources is strongly expected *a priori*, or where the length of the data allows for vanishing leakage effects, the above restriction may be lifted.

##### Implementation for target-frequency specific information transfer

To measure the target-frequency specific *mTE*, we apply the same algorithm as before, but this time we create frequency-specific surrogate data from the target time series (also see Figure S1 supplementary material):

1. Perform a wavelet decomposition through the MODWT to obtain a time-frequency representation of the target time series’ present state, *T_t_*, the target of the multivariate information transfer from **S**_*i*_ to **T**.
2. At the *j^th^* scale of the MODWT decomposition shuffle the wavelet coefficients to destroy information entering the scale in the target or amplitude-phase relations. This step is different from the shuffling in the source algorithm implementation; here, we destroy only the target current value *T_t_* to obtain 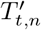, where 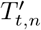 is the *n*-th *target surrogate* under analysis in the network at scale *j* and leaving the target past set, **T**_<*t*_, intact,

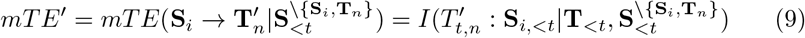
3. Apply the inverse wavelet transform, IMODWT, to reconstruct the time series in the time domain.
4. Compute the *mTE*’ between the source and the *target surrogate,* conditional on all other significant sources in the network.

a. Repeat step 2 to 4 for N permutations to build a surrogate data distribution.
b. Repeat step 1 to 4 for all *J_0_* scales.
5. Check for which scale, *j*, the difference between the original *mTE_tot_* and the median of the *mTE’* distribution is maximal, and determine statistical significance for this scale, similar to the source-frequency implementation.

#### Algorithm II: Testing for direct information transfer from source to receiver frequencies

Consider the following scenario where a certain frequency in the source transfers information to a certain frequency in the target (Fig. 2, A). We would like to then identify these two related frequencies in the source and the target and to determine that there is indeed transfer information *between them*—and not to other, more broadband parts of the spectrum. In other words, we want to exclude the possibility that the source frequency sends information to many other frequencies in the target, potentially even missing the identified target frequency, while the identified target frequency receives information from many source frequencies, potentially excluding the identified source frequency (Fig. 2, B)—such that the direct information transfer between the two identified frequencies is actually absent. We also want to exclude the possibility that the direct information transfer between source and target is entirely redundant with other spectral components of information transfer (Fig. 2, C).

**Fig 2.**
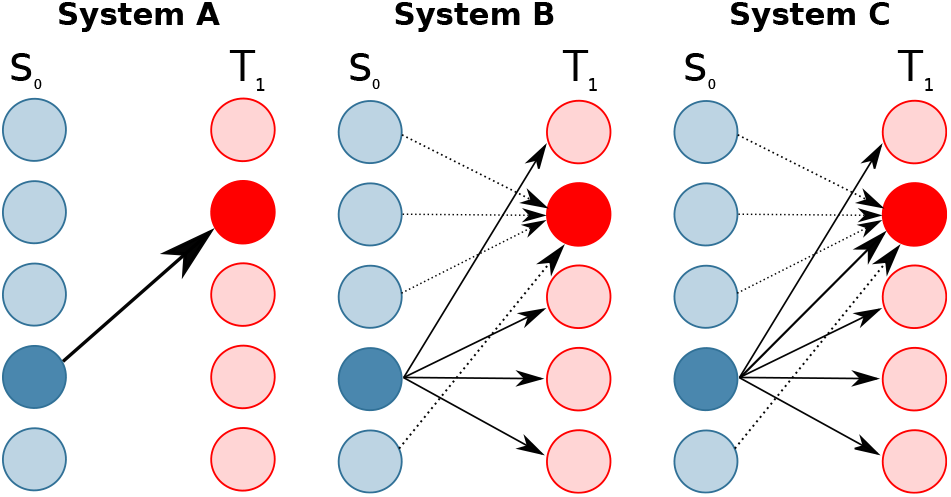
Three systems with the same identified sending and receiving frequencies (indicated by the darker blue and red colors), but a different structure of information transfer. In system A one source and one target frequency take part in a direct transfer of information between them. In system B one source frequency sends information to all target frequencies except the identified target frequency. This one target frequency, in turn, receives other information from all source frequencies except the identified source frequency. In system C the same source frequency sends information redundantly into all target frequencies, while one target frequency receives (partially different) information redundantly from all source frequencies.

When applying algorithm I in the setting assumed above (Fig. 2, A) to the target we will observe a drop in mTE for the surrogate data at the source frequency driving the information transfer, and the target frequency receiving it. If we applied the same algorithm to data where the phase of the sending frequency had been destroyed *beforehand*, then no information transfer should be seen from the source, and thus, also algorithm I applied to the target should also not yield a drop anymore (for the surrogate data with an additionally scrambled target, see also Fig. 3, B and C). Since in this procedure we first swap out the source frequency and then the target frequency in addition, we also refer to algorithm II as the ‘swap-out swap-out (SOSO)’ algorithm from here on. This SOSO algorithm is to be applied after algorithm I has identified specific source and specific target frequencies. That is, we apply the SOSO algorithm as a *post-hoc* analysis.

**Fig 3.**
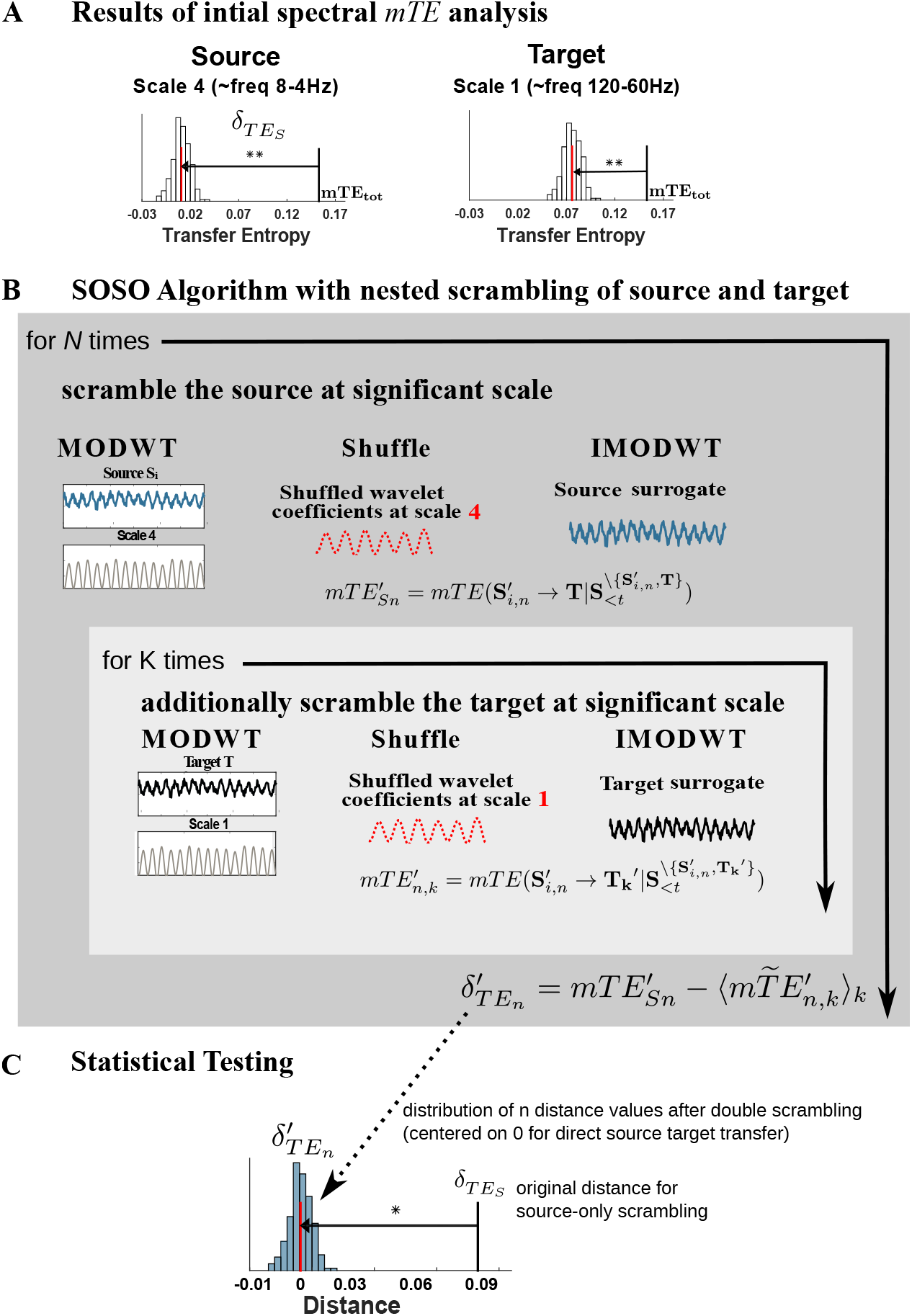
Algorithm II (SOSO). Algorithm to determine whether information transfer exists from an identified information source scale to an identified target scale. (A) Results from the initial analysis using Algorithm I indicating significant information transfer emanating from one scale (here source scale *j* = 4) and significant information reception at a target scale (target scale *r* = 1). (B) To test if the information send from the source scale is indeed the information that is received at the target scale do the following: scramble the source at the relevant scale *N* times and note the 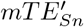 values. For each such scrambled source then apply algorithm I for the target, i.e. scramble the relevant target scale *K* times and note the distribution of the 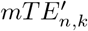 values. Compute the drop in *mTE* obtained for the *n – th* source shuffling with respect to the median of the distribution of source-and-target shuffled 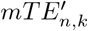 values, 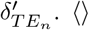-brackets indicate taking the median here. (C) Statistically test the original source drop *δ_TEs_* against the distribution of the 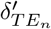. A significantly larger value of *δ_TEs_* indicates that information send by the source scale is indeed received by the target scale.

##### Algorithm implementation

In the following we describe a version of the implementation of the SOSO algorithm, in which we first destroy the source time-series 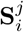 and subsequently also the target **T**^*r*^, where *j, r* are the scales of interest. The algorithm can also be applied in the opposite direction by first destroying the target **T**^*r*^ and subsequently the source 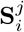.

First, let 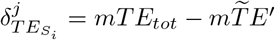, be the distance between the *mTE_tot_* and the 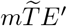, computed with Algorithm I at scale *j* from source **S**_*i*_ and target **T**. Then, Algorithm II comprises two main steps:

1. For *N*-times, scramble the source time-series **S**_*i*_ with one of the implemented shuffling methods and compute the 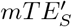 (where the subscript *S* indicate the as described in the subsection 2.2.1 *(Spectral mTE algorithm implementation*). Here the subcribt ‘S’ indicates that the source has been shuffled.

a. For each permutation, create an inner loop running *K*-times, where also the target current value *T_t_* is destroyed at scale *r*, as before, and compute *mTE’*, to obtain a distribution of 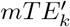 values
b. Compute the distance between 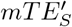 and the median of the distribution 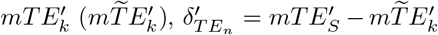 to obtain a distribution of distances
2. Check whether 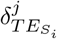 is at the extreme upper end of the surrogates distances distribution 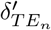.

The SOSO algorithm performs (*N* +1) * *K* mTE computations at the selected source scale and target scale pairs, where *N* and *K* are the number of permutations. Although the SOSO algorithm could be, in principle, applied to all possible source- and target-scale combinations of the identified network, we discourage this approach as pointed out in section *Advantages and drawbacks of the proposed methods.*

### Evaluation

To test the ability of the proposed spectral mTE algorithm to successfully estimate frequency-specific sender-receiver information transfer, we employed multiple synthetic simulations, where the information transfer was known (ground truth). Additionally, we demonstrate the application of algrithms I and II to two neural data sets. The first is a human neuroimaging dataset, acquired with Magnetoencephalography (MEG); the second dataset consists of local field potential (LFP) recordings from the Ferret cortex. The simulations for individual scenarios and details of the neural MEG and LFP data are described below. All analysis were performed with a block permutation of the wavelet coefficients method (to construct surrogates) and LA(8) as mother wavelet, if not stated otherwise.

#### Example 1: Information transfer from one source to one target frequency in a bivariate system

At first, we simulated a simple bivariate scenario, where a single source *S*_0_ is (multiplicatively) coupled to a target *T*_1_ that oscillates at a much faster frequency, such that the amplitude of the target is modulated by the phase of the source, leading to a cross-frequency information transfer (CFIT) (Fig. 4, panel B). Moreover, the source is coupled to the target with a delay of 2 samples. The synthetic data are generated according to the following equations:

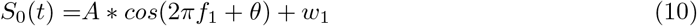

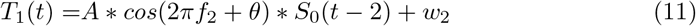

**Fig 4.**
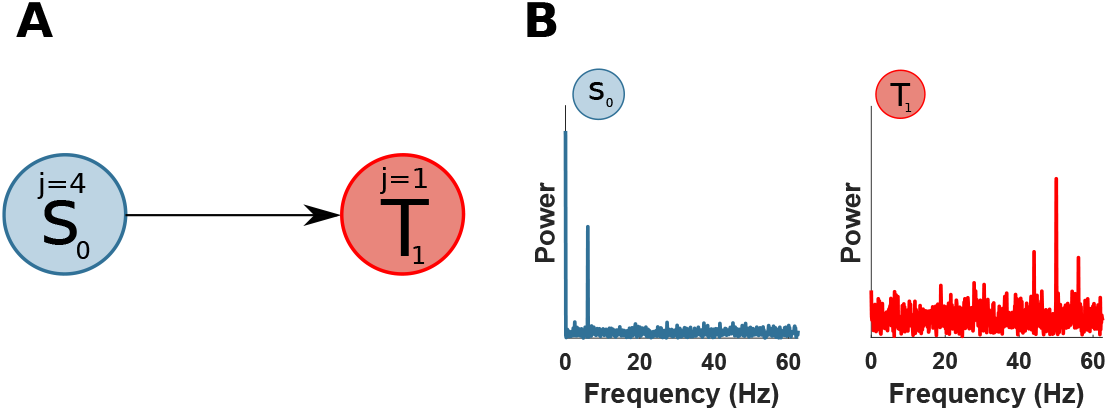
Bivariate simulation with CFIT. (a) A source *S*_0_ is unidirectionally coupled, at scale j=4 (frequency band 4 — 7 Hz), with a target *T*_1_ at scale j=1 (frequency band 31 — 62 Hz). (b) Power spectra of *S*_0_ and *T*_1_.

Where, *A* is the amplitude of the signal and is set to 1 if not stated otherwise, θ is a uniform random variable between 0 and 2*π*, *f*_1_ = 6 Hz, *f*_2_ = 50 Hz and *w*_1_, *w*_2_ are samples of i.i.d Gaussian noise process with a standard deviation of 1. We simulated 10 seconds and 100 trials with a sampling rate of 125 Hz.

First, we performed a TE analysis to recover the source-target information transfer. Table 2 reports the result of the TE analysis. We recovered the true direction of interaction from *S*_0_ to *T*_1_, with a max TE at a lag of 2 samples (16 ms), as simulated. Then, we applied the spectral mTE algorithm to the identified source-target relation to recover the sender- and receiver-frequency information transfer. Fig. 5 shows a significant drop of *mTE*\* in the source *S*_0_ at source-scale 4 (frequency band 4 – 7 Hz), as expected. The amplitude-phase modulation of the target *T*_1_ is visible at target scale 1 (frequency band 62 — 31 Hz)—again as expected. In this relatively simple scenario in terms of sender-receiver frequency relation, the spectral mTE is able to perfectly recover the information transfer in terms of identifying the correct frequencies via the scale of the maximal drop of the surrogate-based distribution and with statistical significance at those frequencies.

**Fig 5.**
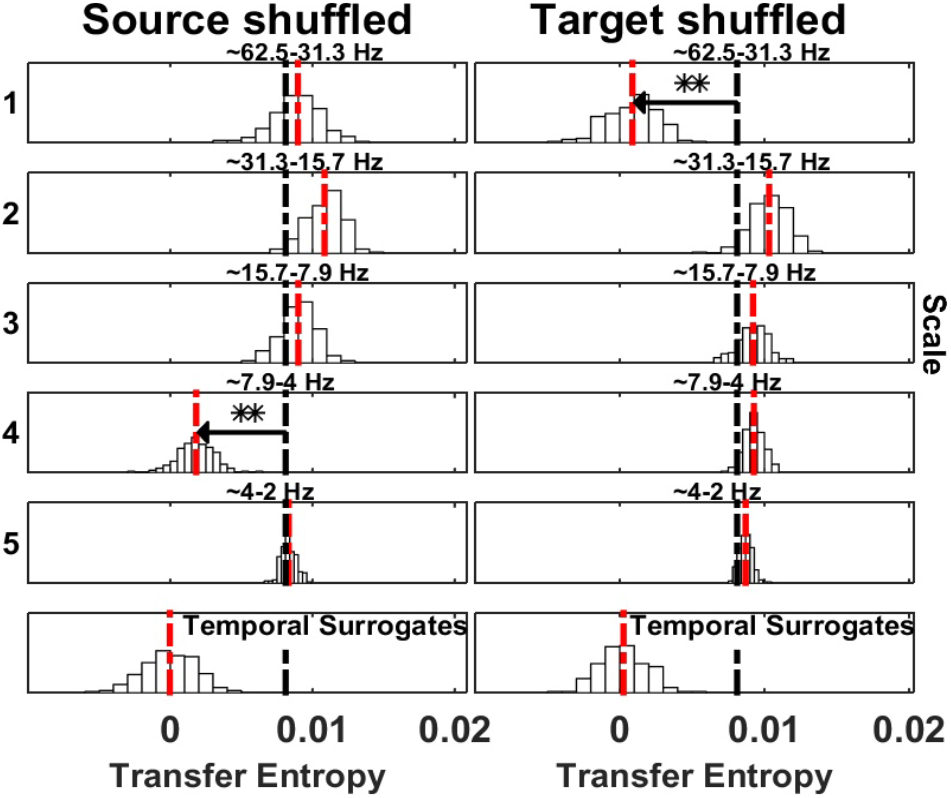
Spectrally resolved Transfer Entropy for example 1. Each panel, except those at the bottom, shows the *mTE*’ distribution obtained from the surrogate datasets with shuffled coefficients at the scale indicated to the left, or, equivalently, the frequency band indicated at the top of each panel. White bars indicate relative frequency, the red dashed line is the median of the surrogate *mTE*’ distribution, the black dashed line is the original *mTE* value. The horizontal black line indicates the distance *δ_TE_* between the original *mTE* and the median of the surrogate distribution (**, *p* < 0.005; *, *p* < 0.05). These display conventions will be kept for figures displaying spectrally resolved TE analyses. Information transfer correctly drops when wavelet coefficients are selectively shuffled at scale 4 at the source (*S*_0_, left column). The corresponding reception of information at the target (*T*_1_) is shown on the right, where a drop for shuffled wavelet coefficients is observed for the frequency band receiving the information in this simulation (i.e. scale 1). The temporal surrogate analysis using surrogates constructed by permuting blocks of samples in the time-domain is shown in the bottom row.

**Table 1.** Results of TE analysis.

| simulation system | interaction $source \rightarrow target$ | TE max lag (ms) | $p$-values |
| --- | --- | --- | --- |
| *Example 1* | $S_0 \rightarrow T_1$ | 16 | $<0.01^{**}$ |
| *Example 2* | $S_0 \rightarrow T_1$ | 4.1 | $<0.01^{**}$ |
| *Example 3* | $S_1 \rightarrow T_0$ | 4 | $<0.01^{**}$ |
| *Example 4* | $S_0 \rightarrow T_1$ | 16 | $<0.01^{**}$ |
| *Example 5* | $S_0 \rightarrow T_1$ | 8 | $<0.01^{**}$ |
| *Example 6* | $S_0 \rightarrow T_1$ | 4 | $<0.01^{**}$ |

\* $p < 0.05$ ; \*\* $p < 0.01$ ; \*\*\* $p < 0.001$ ;

**Table 2.**
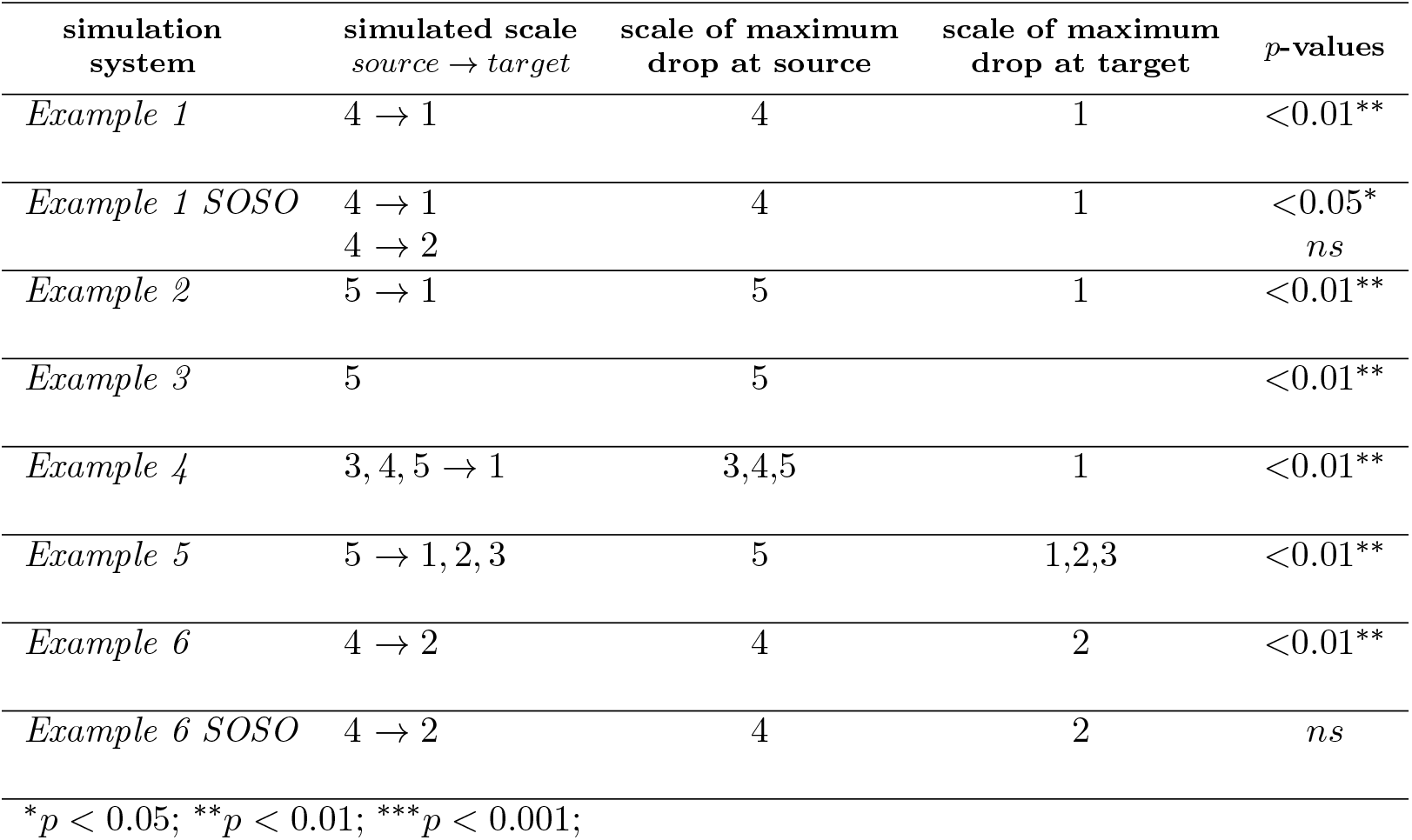
Results of Spectral TE analysis.

#### Evaluation of the SOSO Algorithm (II) on example 1

Here, we evaluated the SOSO algorithm on the CFIT from above (section *Example 1: Information transfer from one source to one target frequency in a bivariate system*). In the first SOSO analysis we set the source scale to be tested to *j* = 4 and the target scale to *j* = 1 - as these were revealed by the spectral mTE analysis. As a control analysis, intended only for demonstration purposes here, the target scale was set to *j* = 3, i.e. a scale not identified as receiving information.

In the first SOSO analysis, the distance 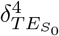 was significantly bigger than the median of the distribution of the 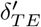 (see Table 2 and Fig. 6 left panel), when we simultaneously destroyed source and target specific scale, indicating a direct information flow between the source and the target. In contrast, and as expected, no significant difference was found when we set the target scale to *j* = 3, since the simulated information transfer between source scale *j* = 4 and target *j* = 1 was not removed by shuffling at the wrong scale (i.e. *j* = 3). In section *Relation of the partial-information framework and the SOSO-algorithm* we further outline the importance of the SOSO algorithm in terms of the PID framework.

**Fig 6.**
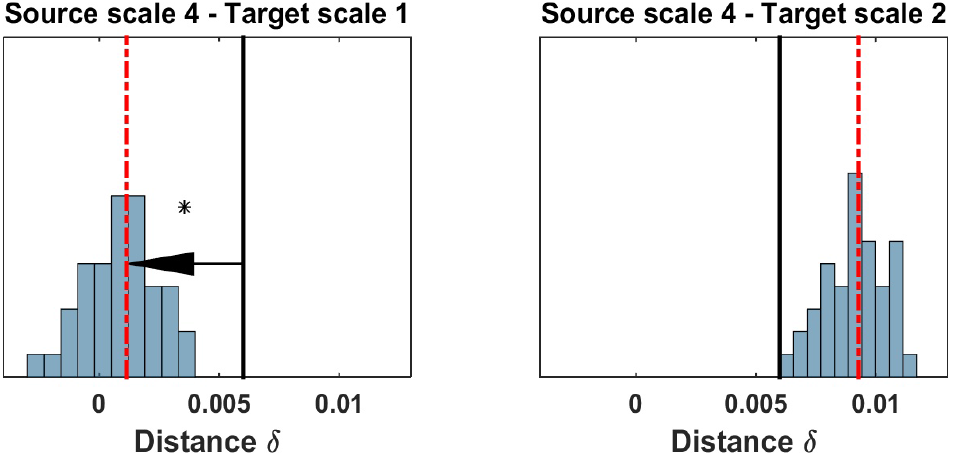
SOSO applications to the bivariate cfit simulation (example 1). Blue bars display the distribution of distances 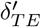 between the median of the surrogate data distribution with a shuffled source (compare Fig. 5) when also the target is shuffled, the red line indicates the median of the distribution of 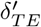, the black line indicates the original distance between the median of the surrogate data distribution with a shuffled source and the *mTE* value computed on the original data. If this latter value is found in the upper rejection interval of the distribution of 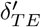, there is significant direct information transfer from the source to the target frequency band under investigation. (Left Panel) No information transfer remains when the source sending scale and the target receiving scale are simultaneously shuffled and no drop of *mTE* can be seen (the distribution 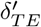 approaches 0); the original drop in *mTE* is significantly larger. (Right panel) Information transfer remains when an unrelated target frequency band is shuffled. 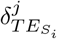 (black bar), median of the 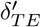 distribution (red dotted bar).

#### Example 2: Cross-frequency information transfer (CFIT) with nonlinear coupling in a multivariate three-node system

Next, we generated a multivariate network of three nodes. The network simulation was generated as follows: first, a source *S*_0_ was coupled to a target with a CFIT. We adapted an example of [27] to simulate a more complex scenario with a non-sinusoidal driver, since in nonlinear systems such as the brain, perfect sinusoidal are often an exception [28]. The CFIT coupling was between *f*_1_ = 6 Hz and *f*_2_ = 80 Hz. To modulate the amplitude of the target time series we employed a sigmoid on the source and a delay of 2 samples. Second, to create a multivariate network, a ‘distractor’ node *S*_2_ was added with an oscillation of *f*_3_ = 90 Hz and it was modulated independently of the *f*_1_ = 6 Hz of *S*_0_ (Fig. 7, panel A). The simulation consisted of 10 seconds and 50 trials with a sampling rate of 240 Hz. The driver *S*_0_(*t*) was generated applying a bandpass filter to a Gaussian white noise at center frequency *f*_1_ with a bandwidth of 0. 4 Hz. The *S*_2_(*t*) and *T*_1_ (*t*) were generated as follow:

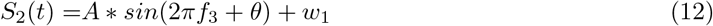

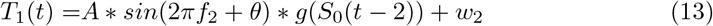

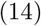

where *g*(*x*) is the sigmoid function:

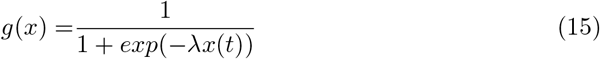

with λ = 3. A Gaussian white noise with a standard deviation of 1.2, 0.8, 0.6 was added to the signal *S*_0_, *S*_2_ and to the target *T*_1_, respectively.

**Fig 7.**
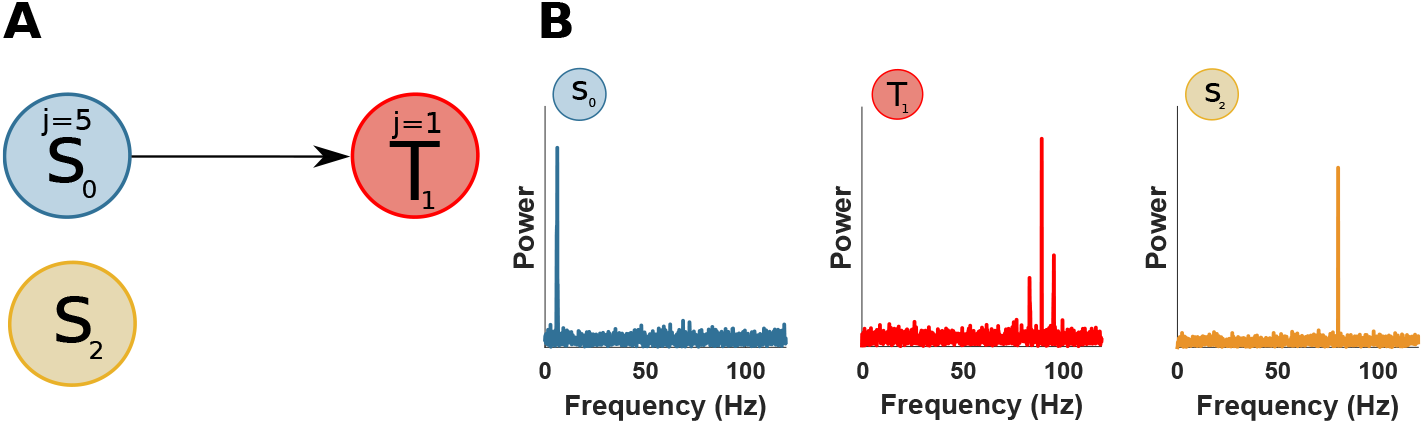
Multivariate simulation with CFIT. (a) A source *S*_0_, but not *S*_2_, is unidirectionally coupled, at scale *j*=5 (frequency band 4-8 Hz), with a target *T*_1_ at scale j=1 (frequency band 60-120 Hz). (b) Power spectral of *S*_0_, *S*_2_ and *T*_1_.

The TE analysis recovered the multivariate network with the associated delay (table 2), identifying significant TE only between *S*_0_ and *T*_1_. The spectral mTE revealed the CFIT between *S*_0_ and *T*_1_, with the maximal distance from the *mTE_tot_* at scale 5 for *S*_0_ and scale 1 for *T*_1_ (Fig. 8).

**Fig 8.**
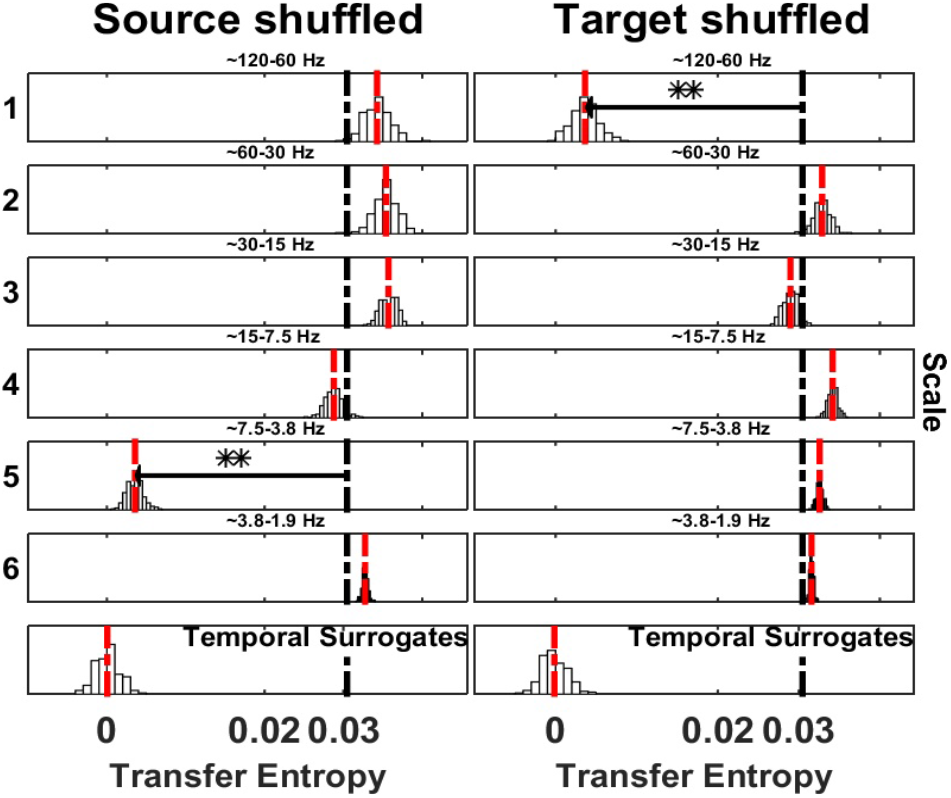
Spectrally resolved Transfer Entropy for example 2. See Fig. 5 for display conventions. (Left panel) Information transfer, correctly, drops when wavelet coefficients are selectively shuffled at scale 5 (frequency band 4-8 Hz) on the source site. The corresponding reception of information at the target is shown on the right panel, where a drop for shuffled wavelet coefficients is observed at scale 1 (frequency band 60-120 Hz), which contained the simulated target frequency.

#### Example 3: Delayed-coupled Rossler systems (nonlinear)

We evaluated the spectral *mTE* with a non-linear system able to generate self-sustained non-periodic oscillations. To this end we generated a coupled Rössler oscillator similar to [29]. The model was simulated with the following equations:

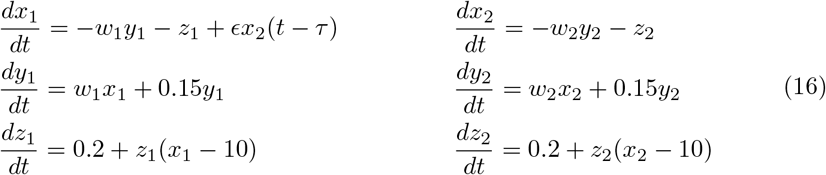

where *w*_1_ and *w*_2_ are the natural frequencies of the oscillator which were set to 0.8 and 0. 9, *ϵ* = 0.07 is the coupling strength and *τ* is the time delay, which was set to 2 time steps. As can be seen in Fig. 9, the two systems oscillated around 8 Hz, but were not identical. The analysis was performed on the assumption that only variables *x*_1_(*t*) and *x*_2_(*t*) could be observed, with *S*_1_ = *x*_2_(*t*) and *T*_0_ = *x*_1_(*t*), in this simulation.

**Fig 9.**
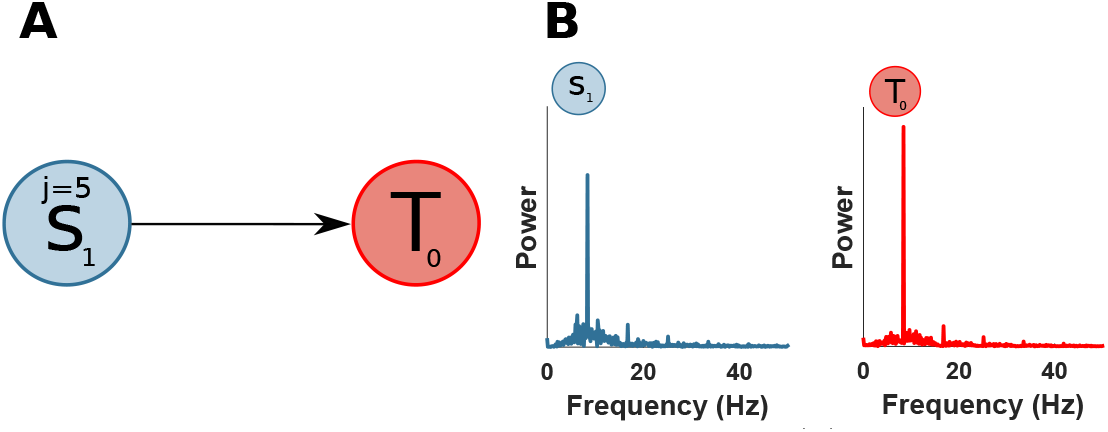
Delay coupled Rossler system simulation. (a) A source *S*_1_ is unidirectionally coupled, at scale j=5 (frequency band 8-16 Hz), with a target *T*_0_. (b) Power spectra of *S*_1_ and *T*_0_.

The TE analysis correctly identified the driver *S*_1_ with a sample delay of 2 (table 2). The spectral mTE showed a significant drop at scale 5 (Fig. 10). No significant drop was observed at the target site *T*_0_.

**Fig 10.**
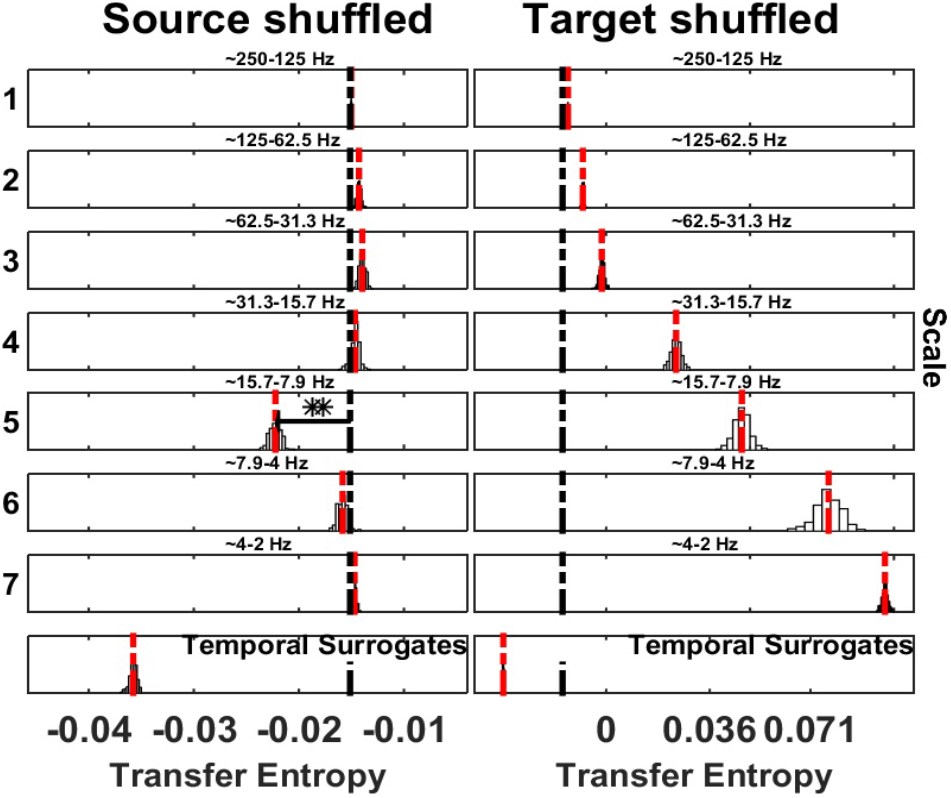
Spectrally resolved Transfer Entropy for example 3. See Fig. 5 for display conventions. (Left panel) Information transfer, correctly, drops when wavelet coefficients are selectively shuffled at scale 5 (frequency band 8-16 Hz) on the source site. (Rigth panel) No significant drop is present at the target site

#### Example 4: Information transfer from multiple source frequencies to one target frequency

To test the ability of the spectral *mTE* to recover multiple source frequencies sending information to a target frequency, we simulated a bivariate example similar to *Example 1* but with multiple source scales showing a phase-amplitude relation with a single target scale.

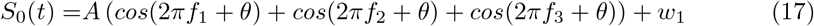

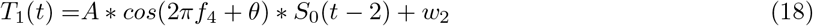

with *S*_0_ constructed as a sum of sinusoids with different phases *θ* and Gaussian noise processes for *w*_1_ and *w*_2_, as before. Then, the source *S*_0_ modulates the amplitude of the target *T*_1_ at scale *j* = 1 with a sample delay of 2 (Fig. 11). The simulation consisted of 5 seconds with 50 trials at 125 Hz.

**Fig 11.**
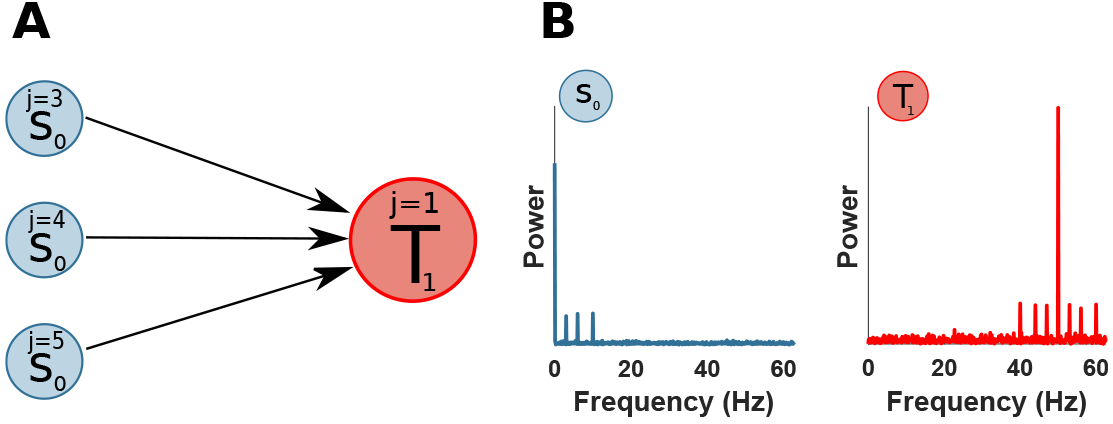
Multiple sending sources bands to a target. (a) A source *S*_0_ is unidirectionally coupled, at multiple scales: j=3 (frequency band 8-16 Hz), j=4 (frequency band 4-8 Hz) and j=5 (frequency band 2-4 Hz), with a target *T*_1_ at scale j=1 (frequency band 31-63 Hz). (b) Power spectral of *S*_0_ and *T*_1_.

The TE analysis showed the source *S*_0_ as driver of the target *T*_1_ (Table 2), as simulated with a sample delay of 2. Then, we applied the Spectral TE to identify the three scales of the source *S*_0_ sending information to the target scale *j* = 1. Fig. 12 showed three significant scales: 3 (frequency band 8-16 Hz), 4 (frequency band 4-8 Hz), 5 (frequency band 2-4 Hz), at source site and scale 1 (frequency band 31-63 Hz) at target site. This application demonstrates the ability of the spectral TE to recover multiple sources sending information to the target site. We also noted that the three scales at the source have a slightly different drop (or maximal distance from original *mTE_tot_*) which can be related to how noise affects the wavelet frequency decomposition at different scales.

**Fig 12.**
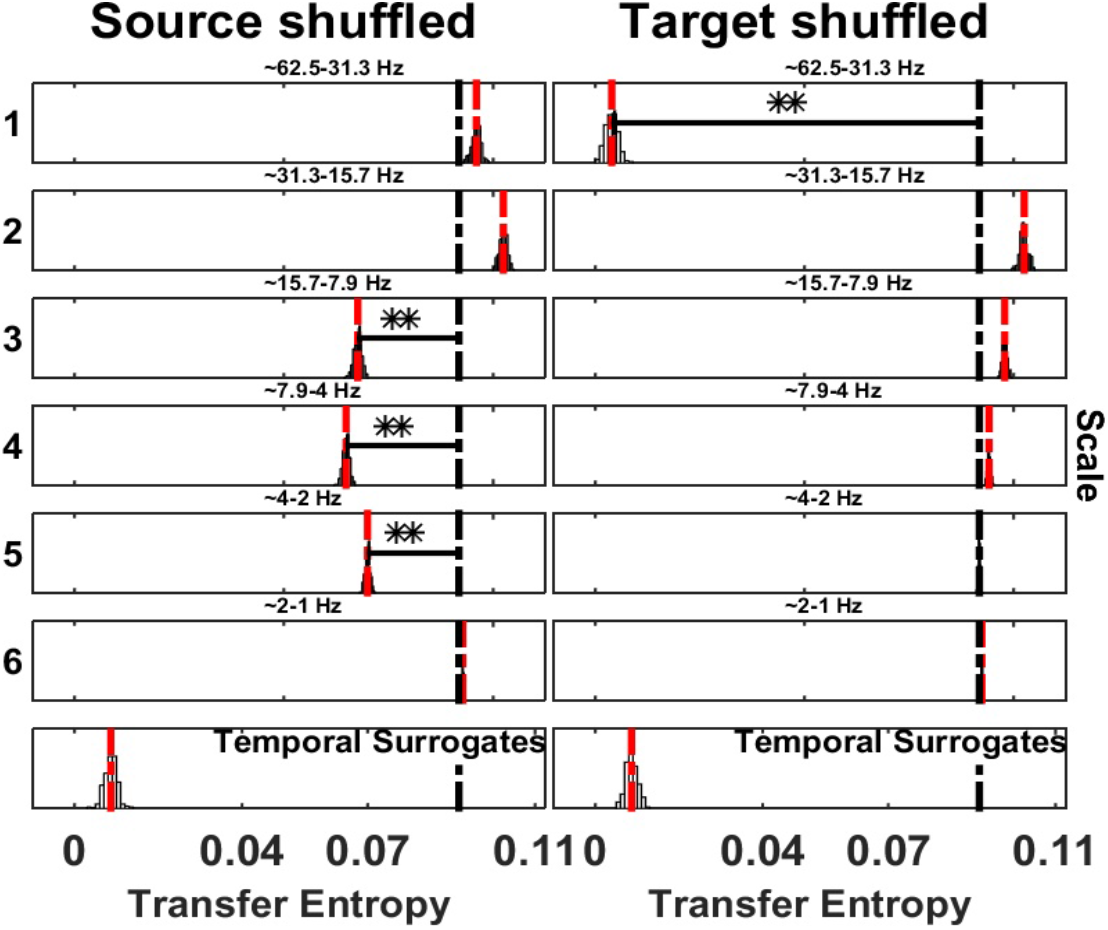
Spectrally resolved Transfer Entropy for Example 4. See Fig. 5 for display conventions. (Left panel) Information transfer, correctly, drops when wavelet coefficients are selectively shuffled at scale 3 (frequency band 8-16 Hz), 4 (frequency band 4-8 Hz), 5 (frequency band 2-4 Hz) on the source site. The corresponding reception of information at the target is shown on the right panel, where a drop for shuffled wavelet coefficients is observed at scale 1 (frequency band 31-63 Hz).

#### Example 5: Information transfer from one source frequency to multiple target frequencies

To test the ability of the spectral *mTE* to recover one source frequency sending information to multiple target frequencies, we simulated a bivariate example with a single source frequency showing a phase-amplitude relation with multiple target frequencies.

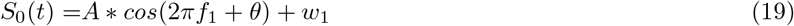

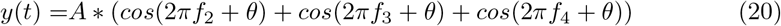

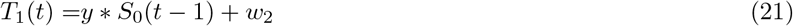

where *y* is a sum of sinusoids at different high frequencies, with *f*_2_ = 100 Hz, *f*_3_ = 58 Hz and *f*_4_ = 30 Hz, which are modulated by the source *S*_0_ with *f*_1_ = 5 Hz, with a sample delay of 1 (Fig. 13), and different noise levels, *w*_1_ with a standard deviation of 0. 4 and *w*_2_ with a standard deviation of 1. The simulation consisted of 5 second with 50 trials at 250 Hz.

**Fig 13.**
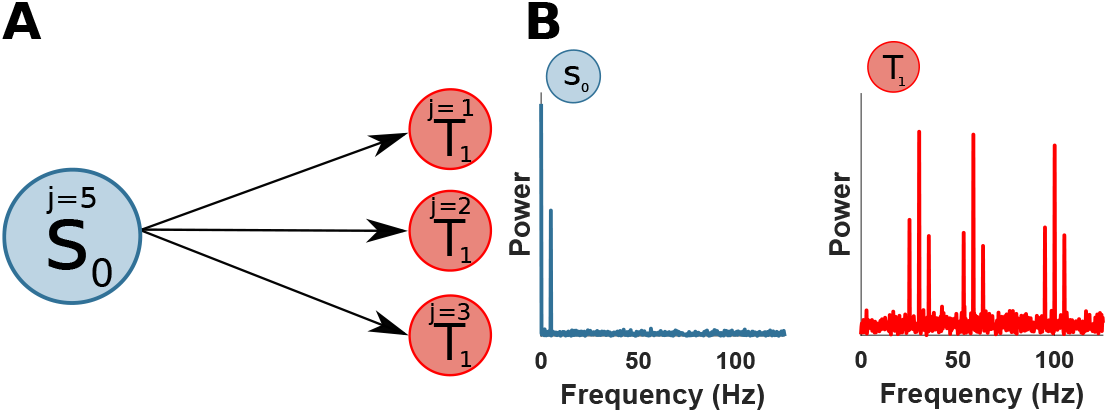
One sending source band to multiple receiving target bands. (a) A source *S*_0_ is unidirectionally coupled, at scales j=5 (frequency band 4-8 Hz), with a target *T*_1_ at multiple scales: j=1 (frequency band 63-125 Hz), j=2 (frequency band 31-63 Hz) and j=3 (frequency band 16-31 Hz). (b) Power spectral of *S*_0_ and *T*_1_

The TE analysis correctly identified the source *S*_0_ as the driver of *T*_1_ with the sample delay of 1 (Table 2). The application of the spectral TE showed two significant scales at the source site: 5 (frequency band 4-8 Hz) and 6 (frequency band 2-4 Hz). In this application, the simulated source scale (*j* = 5) is recovered by the spectral analysis and it is the scale with the largest drop (see Fig. 14). However, scale 6 is also significant. At the target site three scales have a significant drop: 1 (frequency band 63-125 Hz), 2 (frequency band 31-63 Hz) and 3 (frequency band 16-31 Hz), as simulated. This application showed the ability of the spectral TE algorithm to identify multiple receiving targets from one source. Although, the scale with the largest drop, reliably reflects the ground truth, caution needs to be posed to nearby scales, which might have a substantial smaller drop, but stillwenficant, if the the spectral decomposition is not perfectly confined within a single band. To obtain a better frequency concentration, we repeated the analysis employing the MODWT with LA(16). Using a longer wavelet filter should decrease the spectral leakage at nearby scales although increasing the number of boundary-coefficients. Indeed, this analysis revealed a correct identification of the only simulated scale 5 (frequency band 4-8 Hz) at the source site, see *Supplementary Material* (Fig S1).

**Fig 14.**
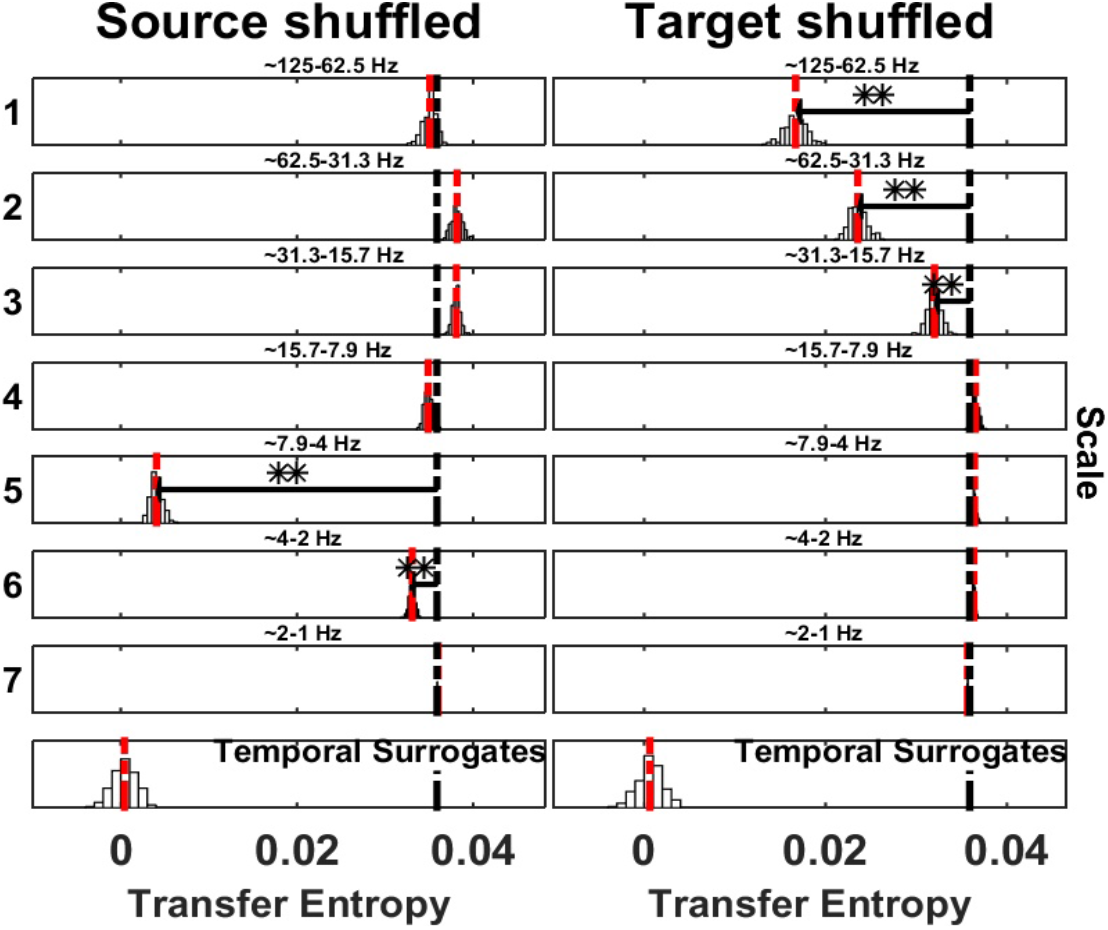
Spectrally resolved Transfer Entropy. See Fig. 5 for display conventions. (Left panel) Information transfer, drops when wavelet coefficients are selectively shuffled at scale 5 (frequency band 4-8 Hz) and 6 (frequency band 2-4 Hz) on the source site. The corresponding reception of information at the target is shown on the right panel, where a drop for shuffled wavelet coefficients is observed at scale 1 (frequency band 63-125 Hz), scale 2 (frequency band 31-63 Hz) and scale 3 (frequency band 16-31 Hz).

#### Example 6: Multiple information flows at multiple frequencies from source to target

In this final simulation, we tested the ability of the SOSO algorithm to rule out direct information flow from a source to a target when one source frequency sends information redundantly into all target frequencies, while one target frequency receives (other) information redundantly from all source frequencies (see Fig. 15 and also Fig. 2). The simulation was carried out according to:

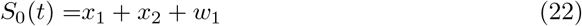

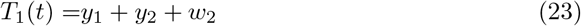

where *x*_1_ and *y*_1_ were simulated with:

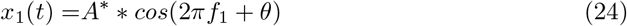

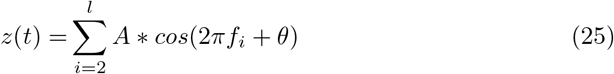

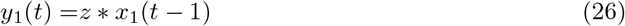

where *l* = 6 and with *f*_1_ = 9 Hz (*j* = 4), *A*\* set to 2, *f*_2_ = 80 Hz (*j* = 1), *f*_3_ = 40 Hz (*j* = 2), *f*_4_ = 18 Hz (*j* = 3), *f*_5_ = 9 Hz (*j* = 4) and *f*_6_ = 5 Hz (*j* = 5). The signals *x*_2_ and *y*_2_ were simulated with:

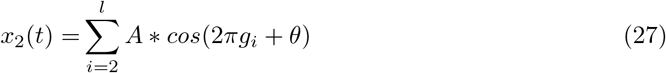

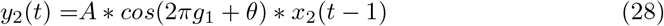

where *l* = 6 and with *g*_1_ = 40 Hz (*j* = 4), *g*_2_ = 80 Hz (*j* = 1), *g*_3_ = 40 Hz (*j* = 2), *g*_4_ = 18 Hz (*j* = 3), *g*_5_ = 9 Hz (*j* = 4) and *g*_6_ = 5 Hz (*j* = 5). The parameter *θ* is a uniform random variable between 0 and 2*π*, *w*_1_,*w*_2_ are samples of i.i.d Gaussian noise process with a standard deviation of 1, and all parameters *A*, except *A*\*, are set to 1.

**Fig 15.**
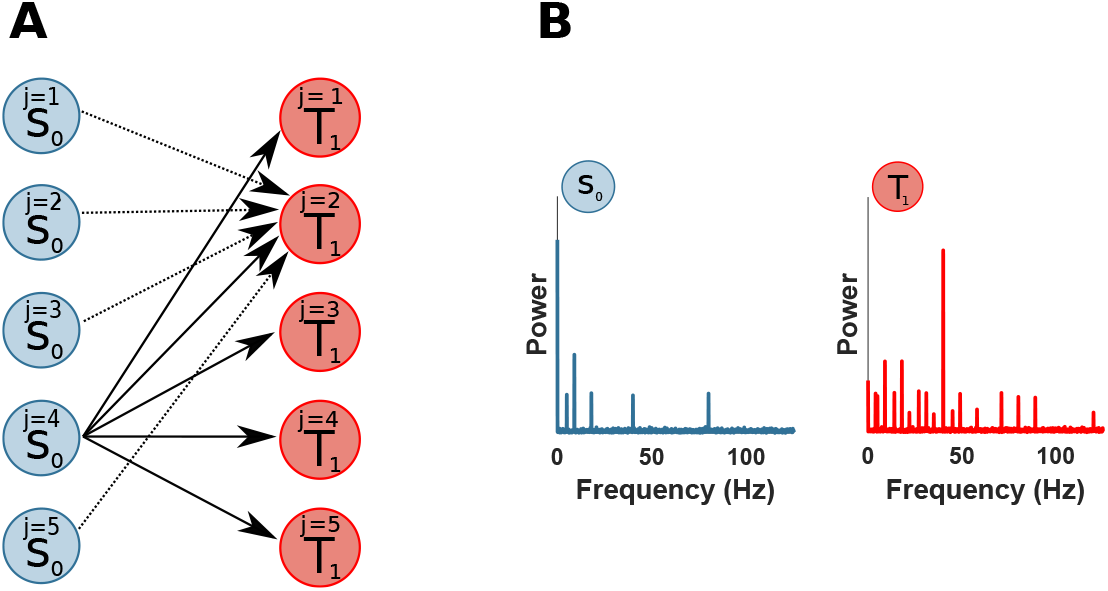
Redundant information flow from source frequencies to target frequencies. (a) A source *S*_0_ is unidirectionally coupled, with target *T*_1_. Multiple scales (*j* = 1, 2, 3, 5) of *S*_0_ are coupled with a single target scale 2 and at the same time a single source scale 4 of *S*_0_ is coupled with multiple target scales (*j* = 1, 2, 3, 4, 5). (b) Power spectrum of *S*_0_ and *T*_1_.

The spectral TE analysis showed, correctly, the largest drop at scale 4 (frequency band 8-16 Hz) at the source site (it was simulated with higher amplitude *A*\* = 2), because this is the source of non-redundant information; additionally scale 5 (frequency band 4-8 Hz) was significant, likely due to spectral leakage. No other scales could be detected at the source site. At the target site, the largest drop was at scale 2 (frequency band 31-63 Hz), being the scale coupled with source scale 4 but also receiving information from all others source scales, i.e. receiving multiple non-redundant information streams. Additionally, only scale 1 (frequency band 63-125 Hz) and 3 (frequency band 16-31 Hz) were significant at the target site. This example was designed to show that a direct non-redundant information transfer from source scale 4 to target scale 2 can be ruled out by using the SOSO-Algorithm. Indeed, the SOSO algorithm confirmed that no direct, non-redudant information transfer took place (see Fig. 17). We direct the reader to section *Relation of the partial-information framework and the SOSO-algorithm*, for further discussion of the SOSO-algorithm in relation to the PID framework.

**Fig 16.**
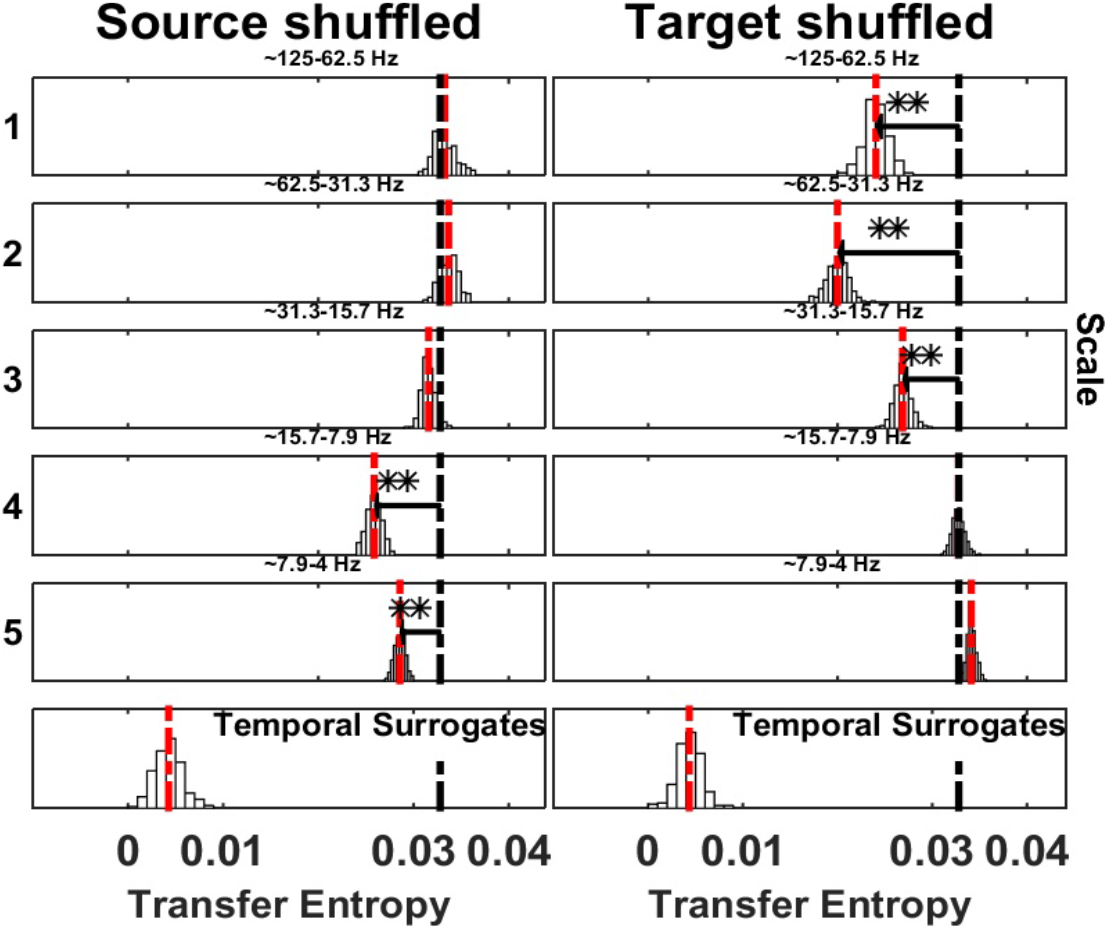
Spectrally resolved Transfer Entropy. See Fig. 5 for display conventions. (Left panel) Information transfer drops when wavelet coefficients are selectively shuffled at scale 4 (frequency band 8-16 Hz) and 5 (frequency band 4-8 Hz) on the source site. The corresponding reception of information at the target is shown on the right panel, where a drop for shuffled wavelet coefficients is observed at scale 1 (frequency band 63-125 Hz), scale 2 (frequency band 31-63 Hz) and scale 3 (frequency band 16-31 Hz).

**Fig 17.**
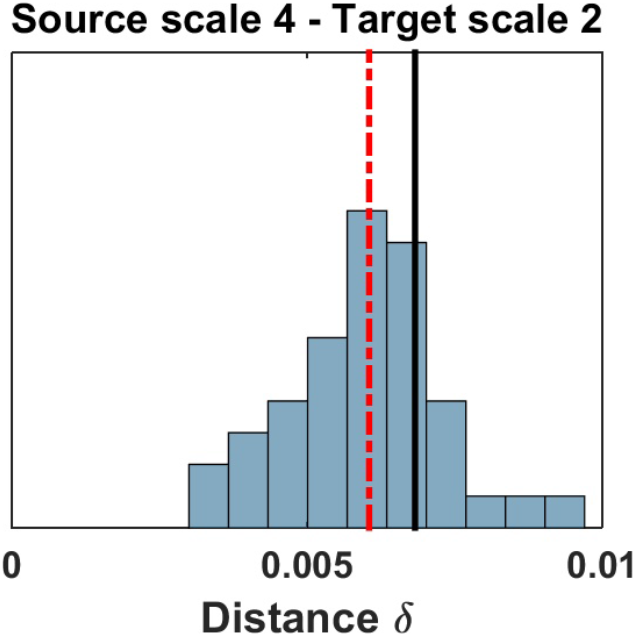
SOSO application to redundant information flow. See Fig. 6 for display conventions. Information transfer remains when the source scale 4 and the target scale 2 are simultaneously shuffled, ruling out a direct information transfer between these two frequency bands.

#### Application to neural data

Finally, we tested the spectral TE method on two different neurophysiological datasets, one human MEG dataset with significant TE between seven sources published in [30], and one local field potential (LFP) recording in the ferret in Prefrontal Cortex (PFC) and Primary Visual Area (V1) published in [31].

##### Information transfer in MEG data from a Mooney face detection task

The data analyzed here were published in [30]. In short, neural activity was recorded with MEG from n=52 subjects at 1.2 kHz sampling rate on a 275 channel whole head magnetoencephalograph (Omega 2005, VSM MedTech). Subjects had to detect either faces (face condition) or houses (house condition) in a stream of black and white pictures of faces (Mooney faces), houses, and scrambled versions of these. From the MEG recordings task-relevant sources were identified by means of beamformer source reconstruction from pre-stimulus baseline data, reflecting the subject’s expectations relevant for the original study, and a comparison of the local active information storage values between the two experimental conditions. After identifying 5 task-relevant sources, bivariate TE was computed on the baseline interval between all pairs of sources and compared between conditions. This procedure identified a significantly different TE between the two conditions (face, house conditions) from anterior inferotemporal cortex (aIT) to the fusiform face area (FFA). Here we follow up on these results by analyzing which source and target frequencies carried the TE found in the original study.

At the group level, for each scale in the face condition, we determined (with a dependent samples t-test, Monte Carlo estimate [32]) if the 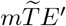 was significantly smaller than the *mTE_tot_* (alpha level was set at 0.05 and Bonferroni corrected for the fourteen scales tested).

In the face condition (Fig 18), we found the aIT source to be sending information at frequencies around 110 Hz (scale 3: 75-150 Hz, with possible additional contributions above 150 Hz scale 2: 150-300 Hz), while the FFA target received significant information transfer at multiple high frequency bands: above 150Hz (scale 2: 150-300 Hz) and around 110 Hz (scale 3: 75-150 Hz) and multiple lower bands: around 14 Hz (scale 6: 9-19 Hz) and around 7 Hz (scale 7: 5-9 Hz). Results for the spectrally resolved mTE in the house condition were qualitatively similar (data not shown).

**Fig 18.**
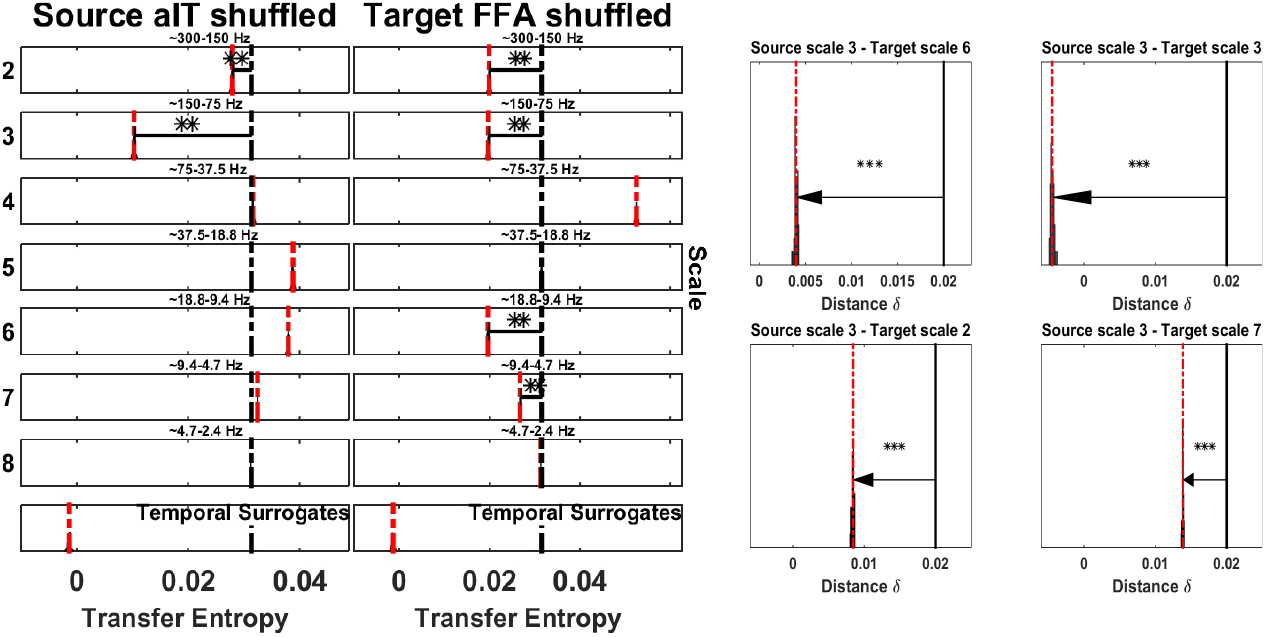
Spectrally resolved information transfer between MEG sources when preparing to detect faces (first panel on the left side). See Fig. 5 for display conventions. Spectrally resolved information transfer between aIT as a source and FFA as a target in the condition where subjects are trying to detect target faces. aIT sends information mainly at 75-150Hz (left column of the first panel), whereas FFA receives information at high frequencies (75-150Hz and above) as well as low frequencies (9-19Hz and 5-9Hz) (right column of the first panel). Analyses of cross-frequency information transfer between specific source frequency in aIT and multiple target-frequencies in FFA (second panel on the right side). All four scales (2, 3, 6, 7) at the target side showed a significant direct information transfer from the source at scale 3. 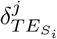 (black bar), median of the 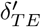 distribution (red dotted bar).

Finally, we applied the SOSO algorithm to the face condition at scale 2 for the source aIT and to the four significant scales at the target FFA (scales: 2,3,6 and 7). The SOSO algorithm showed a significant direct information flow between source scale 2 (frequency band 75-150 Hz) and all four scales at the target site (scales: 2,3,6 and 7), confirming a high complexity of interaction between the source aIT and multiple frequencies at the FFA area (see Fig. 18).

These spectral mTE results provide important information about the spectral complexity of the interaction between aIT and FFA in that the information transfer took place between different frequency bands. These results could not have been obtained with a spectral GC approach, as spectral GC only searches for within-band transfer of information.

##### Occipito-frontal and fronto-occiptal information transfer in the Ferret

We applied the spectral TE method to data from a previous study on anaesthesia effects in the ferret [31]. In short local field potentials were recorded simultaneously in primary visual cortex (V1) and the prefrontal cortex (PFC) of two female ferrets (see Fig. 19 for a schematic depiction of recording sites) under different concentrations of the anesthetic isoflurane and under awake conditions. Since the application of spectrally resolved mTE here served only as a proof of principle we only analysed the data of ferret 1, i.e. the ferret that showed significant bi-directional TE in the awake condition. Moreover, we only analyzed data from the awake condition. For more information on the experimental procedures see [31]. First, using the mTE-algorithms from our new IDTxl toolbox we replicated the earlier findings of a significant bidirectional TE between PFC and V1 and vice versa (see Table 3, and Figs. 20, 22).

**Fig 19.**
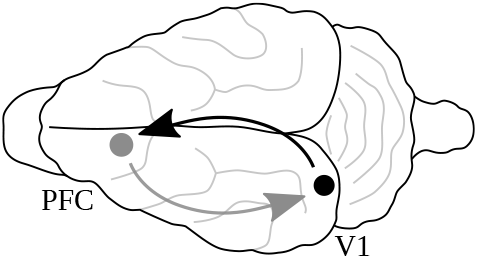
Schematic location of recording sites on the Ferret brain (from [31]).

**Table 3.** Results of TE analysis neural data.

| experiment | interaction | significant subjects | p-values |
| --- | --- | --- | --- |
| *MEG face condition* | *aIT* → *FFA* | 29 | <0.01** |
| *LFP Ferret* | *PFC* → *V1* | n/a | <0.05* |
| | *V1* → *PFC* | n/a | <0.05* |

\* $p < 0.05$ ; \*\* $p < 0.01$ ; \*\*\* $p < 0.001$ ;

Second, in the direction from PFC to V1 the spectral TE method revealed a significant effect in scales 7, 8, 9, with scale 7 (4-8 Hz) as minimum, when PFC was considered as source of V1. In contrast, at the target site (V1), the only scale that revealed a significant TE decrease when shuffled was scale 2 (125-250 Hz, high gamma band, or very high frequency oscillations, VHF), indicating a possible CFIT from PFC to V1, results are reported in table 4. To test this, we applied the SOSO algorithm on scale 7 from the PFC source and scale 2 from the target V1. As a control analysis we additionally randomly picked another scale on the target side to verify that the interaction was restricted only to target scale 2.

**Table 4.** Results of Spectral TE analysis.

| experiment | scale of maximum drop at source | scale of maximum drop at target | p-values |
| --- | --- | --- | --- |
| *MEG faces condition* | 3 | 2,3,6,7 | <0.01** |
| *SOSO* |  |  |  |
| *aIT* → *FFA* | 3 | 2,3,6,7 | <0.001*** |
| *LFP Ferret* |  |  |  |
| *PFC* → *V1* | 7 | 2 | <0.05* |
| *V1* → *PFC* | 3 | 2 | <0.05* |
| *SOSO* |  |  |  |
| *PFC* → *V1* | 7 | 2 | <0.05* |
|  | 7 | 5 | ns |
| *V1* → *PFC* | 3 | 2 | <0.05* |
|  | 4 | 2 | ns |

\* $p < 0.05$ ; \*\* $p < 0.01$ ; \*\*\* $p < 0.001$ ;

Similarly to our simulations (see Fig. 6), the PFC source at scale 7 and the V1 target at scale 2 showed a significant decrease of the distribution of delta values and *δ_TE_S_i___* was in the extreme 5% of the distribution (*p*<0.05*). No significant result was found when we applied the SOSO algorithm to the randomly chosen control target-scale 5 (see Fig. 21 and table 4).

**Fig 20.**
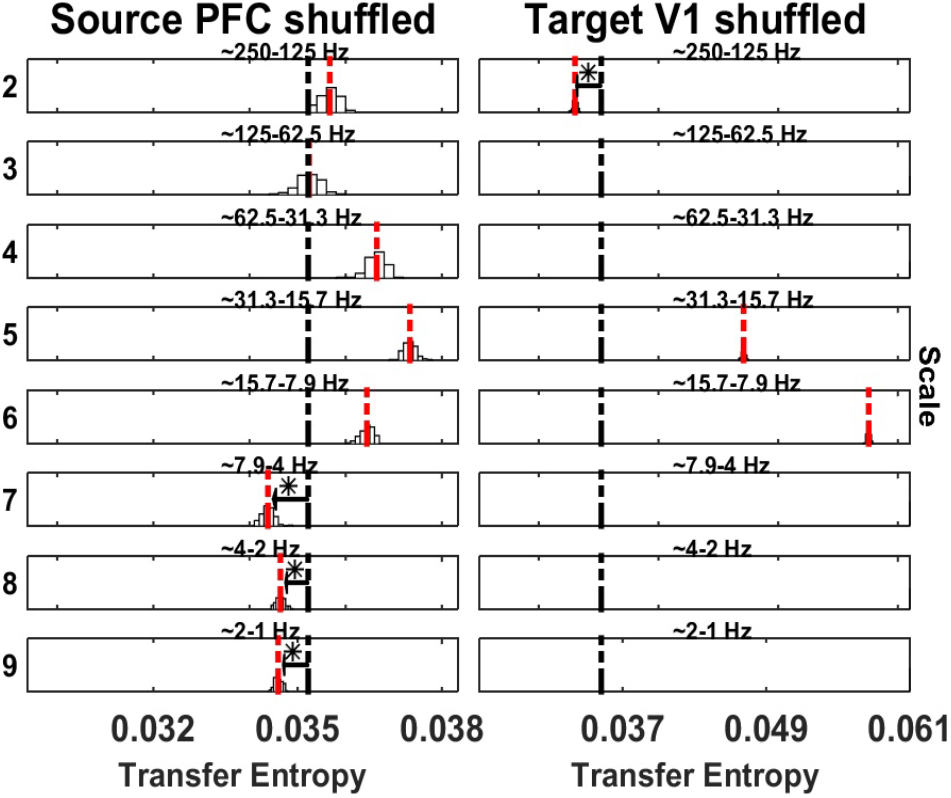
Spectrally resolved information transfer from PFC to V1 in the Ferret. See Fig. 5 for display conventions. (Left panel) Information transfer, drops at scale 7, 8, and 9 on the source site (PFC), when the wavelet coefficients are shuffled. (Right panel) A significant drop is observed at scale 2 at the target site (V1). *mTE_tot_* original (black dotted bar), 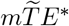 (red dotted bar). Scale 1 is not shown since LFP were low passed at 300 Hz. Temporal surrogates are not shown

**Fig 21.**
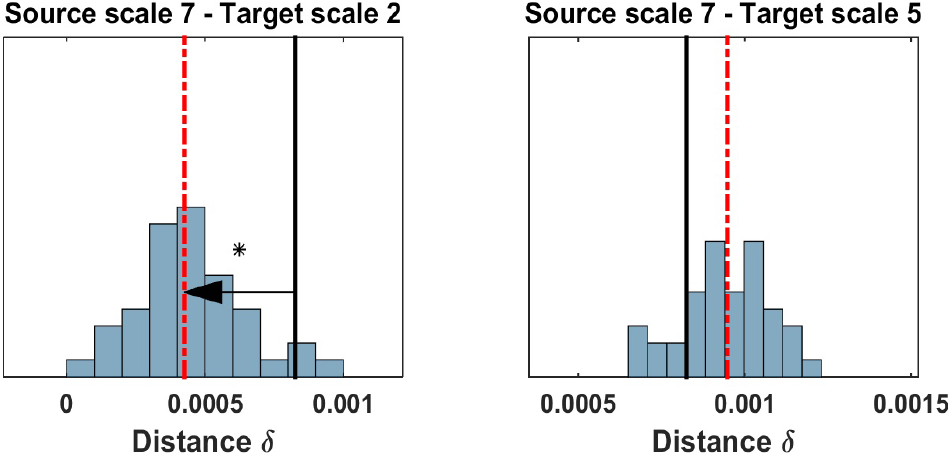
Analyses of cross-frequency information transfer between specific source- and target-frequencies in PFC and V1 of the Ferret. See Fig. 6 for display conventions. (Left Panel) No information transfer is present when the source sending scale and the target receiving scale are simultaneously shuffled and no drop of *mTE* can be seen (the distribution 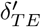 approaches 0). (Right panel) Information transfer is still present when unrelated target frequency band is shuffled. 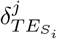 (black bar), median of the 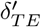 distribution (red dotted bar).

Third, we considered V1 as source and PFC as target. The spectral TE algorithm revealed a significant TE decrease on scale 2, 3, 4 at source site, with scale 3 (high gamma) as minimum. On the target side (PFC) scale 2 was the only significant result (see table 4).

We applied the SOSO algorithm on the source scale 3 and 4, and target scale 2. This analysis revealed a significant decrease of the distribution of delta values for source scale 3 and target scale 2 (*p*<0.05*, see Table 4 and Fig. 23, right panel), but interestingly not for source scale 4 (see Table 4 and Fig. 23, left panel).

**Fig 22.**
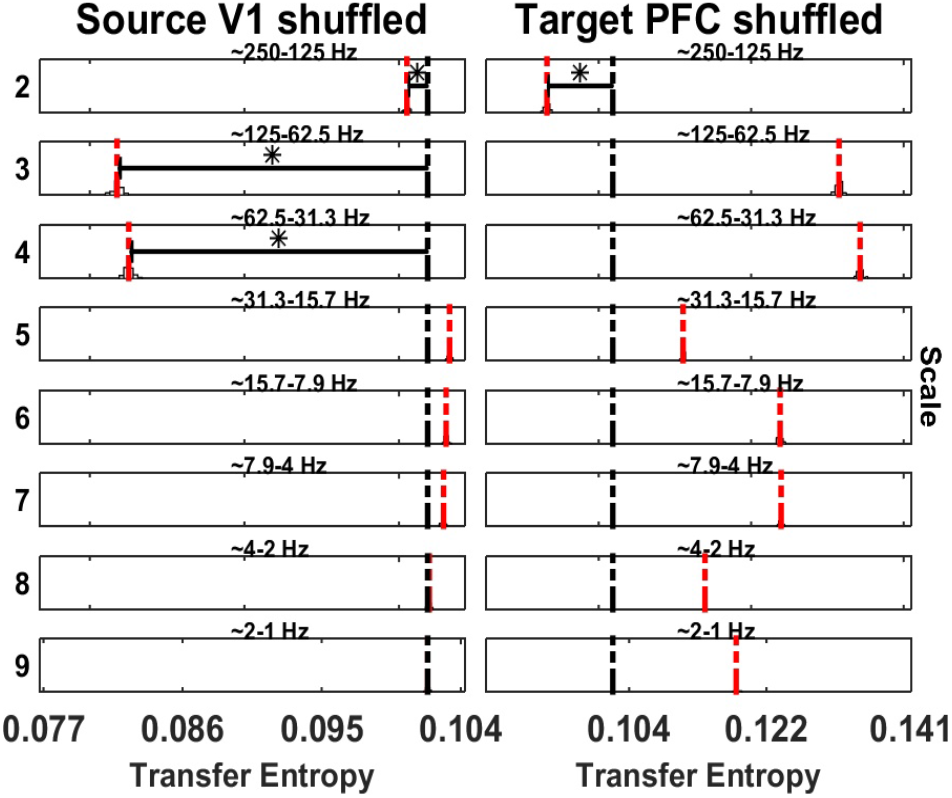
Analysis of spectrally resolved information transfer in the Ferret from V1 to PFC. See Fig. 5 for display conventions. (Left panel) Information transfer, drops at scale 2, 3, and 4 on the source site (V1), when the wavelet coefficients are shuffled. (Right panel) A significant drop is observed at scale 2 at the target site (PFC). *mTE_tot_* original (black dotted bar), 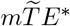 (red dotted bar). Scale 1 is not shown since LFP were low passed at 300 Hz. Temporal surrogates are not shown.

**Fig 23.**
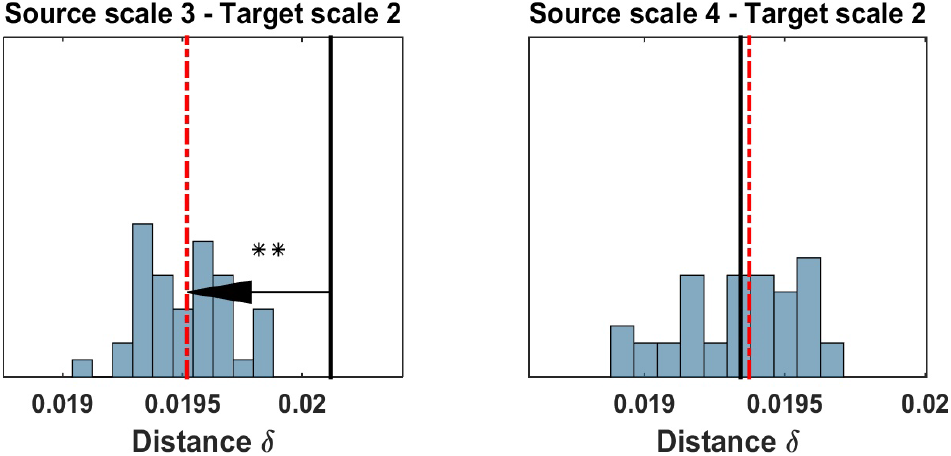
SOSO application to source V1 and target PFC of Ferret. See Fig. 6 for display conventions. (Left Panel) No information transfer is present when the source sending scale and the target receiving scale are simultaneously shuffled and no drop of *mTE* can be seen (the distribution 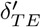 is centered on 0). This means that there is indeed a direct transfer of information from source scale 3 to target scale 2 (Right panel) Information transfer into target scale 2 is still present when the source scale 4 is shuffled, meaning information does not flow from source scale 4 to target scale 2. 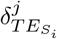 (black bar), median of the 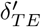 distribution (red dotted bar).

The application of the new spectral TE algorithm on data that showed significant bidirectional TE revealed, a low frequency top-down communication, PFC → *V*1, with a possible CFIT (Fig. 21), and high frequency bottom-up communication V1 → *PFC* (Fig. 22), in agreement with [33].

These results demonstrate the value of separate analyses for source and target frequencies and the post-hoc tests to identify only direct TE from source to target.

## Discussion

We present an algorithm to measure the information transfer that is associated with specific spectral components in an information source or target. Our evaluation results demonstrate that spectral components in the source or target can be reliably identified, given that there are not many closely overlapping components contributing to the information transfer. If many, closely overlapping components are present, a conservative approach is to focus only on the component yielding the largest contribution. One of the advantages of the algorithm presented here is that the original signals are never filtered before the computation of information transfer. Rather, we defer the filtering operation to the creation of spectrally-specific surrogate data, where phase shifts, filtering artefacts or filter-inefficiencies lead to overly conservative results in the worst case, but not to spurious false positives. Our algorithm can be extended to investigate whether the information transferred from a specific source-frequency indeed arrives at a specific target frequency whenever there is an *a priori* reason to assume that this possibility exists in the system under investigation. If no such *a priori* consideration applies, the intricacies of the partial information decomposition (see next section below) warrant utmost caution when inferring a direct transfer of information from identified source to target frequencies.

In the remainder of this section we will detail the added value of spectrally-resolved TE analysis for neuroscience, discuss the inherent but often-overlooked complexities of measuring spectrally-resolved information transfer, discuss advantages and drawbacks of the algorithm presented, discuss its relation to previous approaches, and remark on the choice for the free parameters of the algorithm.

### Information transfer in rhythmic neural processes beyond within-band synchronization and linear interactions

Having the possibility to analyze how information is transferred in relation to various spectral components is of particular interest in neuroscience. This is because of the importance of information transfer for the distributed computation performed in neural systems and because of the prevalence of rhythmic processing in neural systems. This rhythmic processing is evident for example in the mammalian neocortex, where researchers have studied processes in various frequencies for decades (e.g. *δ*, *θ*, *α*, *β*, low and high frequency *γ* rhythms). Our novel analysis techniques offer the unique opportunity to unravel how information is exchanged between different rhythmic processes, but also, and importantly between arhythmic (wideband) and rhythmic processes (see example 4). We thus expect that our methods will widen the current focus on synchronization and within-band interactions to reveal a fuller picture of neural processing.

In this respect, even our proof of concept analysis of MEG and LFP data have provided intriguing new insights:

1. In the MEG data we found two surprising results: (a) Information transfer at very high frequencies that was possibly linked to leaked multi-unit activity or to oscillatory components. We are not aware of other reports of functional connectivity or information transfer at those frequencies, possibly owing to the fact that coupling at these frequency bands may be nonlinear and may not be carried by relatively stable oscillations. Nevertheless our analyses demonstrate that the information in these bands is well captured by the MEG. (b) The information sent via high frequencies from aIT cortex to FFA is also received at the low beta band, i.e. there is information flowing from high frequencies to lower frequencies. This result differs from the usual assumption that lower frequencies modulate higher ones (e.g. via phase-amplitude coupling [34]), but corroborates earlier findings in nonhuman primates that showed the same effect [3].
2. In the Ferret LFP-recordings we observed information being sent from frontal cortices at low frequencies from *δ*, *θ* and *α*-bands, and being received in V1 at high *γ*-frequencies—in line with previous reports. Yet, of the information in the low source frequencies, only information in the alpha band seems to be directly received by the high *γ*-band in the target—with the information sent by the *δ* and *θ*-bands being either redundant with the information in the *α*-band or being received across all frequencies in the target as indicated by the SOSO-analysis (Fig. 21).

In sum, these two exemplary applications to neural data demonstrate the enormous level of detail provided by the proposed algorithms for the analyses of neural communication. These examples also point to the possibility that there are many neural communication processes or mechanisms that have been overlooked so far due to the lack of proper analysis methods.

### Frequency resolved TE as a partial information decomposition problem

To discuss frequency-resolved analysis of information transfer as a partial information decomposition (PID) problem, we first introduce the concept of PID by a simple example. Imagine two source variables *S*_1_, *S*_2_ that together provide some information about a target variable *T*, i.e. the joint mutual information *I*(*T*: *S*_1_, *S*_2_) is non-zero. One may ask, then, how much of that joint mutual information about *T* can only be obtained from *S*_1_, but not from *S*_2_ and vice versa, how much information can be redundantly obtained from either variable, and how much information can be only obtained from *S*_1_, *S*_2_ considered jointly. These three ‘types’ of information are called unique, shared and synergistic mutual information.

In the same way that a joint mutual information can be decomposed we can also decompose a conditional mutual information, e.g. *I*(*T*: *S*_1_, *S*_2_|*Z*). Since transfer entropy is just a conditional mutual information, and since our spectral components would take the role of the source variables *S*_1_,…, *S_n_* we see that asking for the contribution of each spectral component to the overall transfer entropy amounts to solving the partial information decomposition problem. Unfortunately, the full complexity of such partial-information decompositions has only been realized very recently, and their mathematical formulation is still a matter of active research (see for example the recent special issue on this topic [35]), especially when more than two source variables are involved, as will be the norm for a spectral analysis of information transfer.

Moreover, current PID measures only allow a decomposition of either the source or the receiver processes into PID components. This unsolved problem is also the fundamental reason why we have mostly confined ourselves to considering source and receiver frequencies separately (apart from trying to identify the special case of one source frequency interacting with one target frequency). While the field of PID is still under rapid development in terms of proper information theoretic measures, the underlying structure of the problem is clear, and can be harnessed to understand the spectral analysis of information transfer. In particular, using the PID formalism we can clearly map out which specific components and combinations of components of a PID will be detected or missed by our analysis method, irrespective of any particular definition or measure of PID components:

1. Frequencies on the source or target side that contribute unique information to the transfer entropy from source to target will be detected, both on the source and the target side.
2. Frequencies on the source or target side that carry information redundantly (and with approximately equal amount and signal to noise ratio) will be *missed*. This is because destroying one of these frequencies in the surrogate data will not remove the information, as it is redundantly carried via the other frequency. Hence, the mTE on the permuted data will most likely not drop significantly.
3. Frequencies on the source or target side that synergistically contribute to the TE will all be detected, as destroying each of them will reduce the amount of information transfer in the permuted data. There will be no indication as to whether the information transfer was a synergistic or unique contribution of those frequencies, unless there is only one frequency that leads to a reduction of the TE in the permuted data (in which case it is a unique contribution).

We note that the difficulties mentioned under 2. and 3. are generic and do not apply to the analysis of transfer entropy alone but to any spectrally-resolved measure of information transfer. They simply reflect the possibly complex nature of statistical dependencies in multi-variable systems.

### Relation of the partial-information decomposition framework and the SOSO-algorithm

Understanding that measuring frequency-resolved TE is a PID-problem is particularly useful in understanding the necessity of the SOSO-algorithm to determine putative cross-frequency effects (see *Algorithm II: Testing for direct information transfer from source to receiver frequencies*). For system A in Fig. 2 there is direct information transfer from one source to one target frequency. Algorithm I will identify these frequencies and the SOSO-algorithm will confirm the direct transfer of information between these frequencies. In system B, however, one source frequency sends information to all target frequencies except one. This one target frequency, in turn, receives other information from all source frequencies except the identified one. In this system the information sent redundantly by multiple frequencies, and the other information received redundantly will not be revealed, as destroying individual frequencies does not destroy it (information is present redundantly in other frequencies). Yet, the SOSO-analysis will show that the identified source and target frequency do not exchange information directly. In system C the same source frequency sends information redundantly into all target frequencies, while one target frequency receives information redundantly from all source frequencies. Depending on the signal to noise ratio, here either no source and no target frequency will be identified by algorithm I, as all the information is carried redundantly, or both the source and the target frequency will be detected if they have superior signal to noise ratio. Then the direct transfer from one source to one target frequency will be confirmed by the SOSO-algorithm.

Again the difficulties that arise in the identification of sender and receiver frequencies and in establishing a direct transfer of information from one source frequency to one target frequency are closely related to the fact that we deal with a partial information decomposition problem here.

### Methodological advantages and drawbacks of the proposed method

The most important advantage of obtaining frequency-selectivity without filtering the original data is that we do not introduce distortions into the relative timing of the original data or have to deal with other filtering-related artifacts, or even ineffective filters as they were described before. Ultimately this protects us from generating false positive results due to filtering.

With respect to the influence of filtering artifacts biasing TE estimation, as described in [5], manipulating only the surrogate data in the spectral domain restricts the appearance of filtering-related artifacts to the surrogate data. If these artifacts will produce an erroneous increase in TE then this will only lead to conservative errors; if artifacts produce a decrease, this is the desired effect in surrogate data creation, anyways. We consider these filtering-related errors to be mild in most cases, as demonstrated by the correct recovery of relevant source and target frequencies of information flows in our evaluation examples.

Our approach also solves the problem of filter ineffectiveness as described in [7]. This problem arises in approaches that use filtering of the original data to remove information transfer in frequencies of no interest. Yet, filtering will only dampen spectral power, not remove the information contained in a specific frequency as long as the numeric resolution is high enough to keep the unwanted signal above the numeric quantization noise. Ironically, this problem is more serious when using high-precision math libraries. In our approach we use the filtering-equivalent MODWT transform only to *isolate* the information of interest—to then destroy it by coefficient scrambling in surrogate data creation.

In terms of drawbacks, the most important one is likely to be the computational burden our analyses presents, especially when using the SOSO-algorithm. This burden stems from the use of information theoretic estimators for continuous data as well as from recursively nested non-parametric statistics with sufficient iterations and permutations. Due to the computational burden, our method does not lend itself to large-scale exploratory studies as they have been popular with simpler methods based on correlations for example. We therefore advise to apply the methods presented here in a confirmatory way, e.g. for testing highly specific hypotheses of interest, or to neural interactions that have been carefully pre-selected by other methods (e.g. by an mTE analysis, or simply by drawing on prior knowledge).

On the other hand, the hierarchical approach of first searching for source and target frequencies of interest—and only then applying the SOSO-algorithm to selected frequency pairs in a confirmatory step—makes our methods scale better with the number of frequencies involved (basically *O*(*j*) instead of 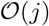, where *j* is the number of frequencies involved).

### Specific caveats

The advantage of the proposed algorithms of avoiding filtering of the original data comes at the price that the measurement of the frequency-specific TE contribution is not in absolute terms, but as a difference to the TE value obtained from the surrogate data. This will lead to a potential underestimation of the information transferred by a certain frequency. Yet, for finite data, a comparison of estimated TE values to the TE obtained from suitable, case-specific surrogate data is recommended in most cases anyway, due to the considerable bias problems inherent in the estimation of information theoretic quantities from finite data [36]. Moreover giving exact quantities for the information sent or received by a frequency again means having to face questions of how to attribute uniquely, redundantly or synergistically sent or received information.

### Relation to previous approaches

While we have seen attempts at frequency resolved TE and mTE estimation at conferences, none of these seem to have been published. The literature on frequency-resolved TE is thus very sparse. Specifically, we found that the approaches in [3,37–39] all use spectral decomposition techniques on the original data, and thus seem suffer from the vulnerabilities detailed in [5,7]. As an exception, Xu and colleagues [40] use a technique similar to the definition of spectral Granger causality, but rely on Gaussianity of the data. Last, we note that the Phase transfer entropy introduced by Lobier [39] as the transfer entropy between the time series of the instantaneous signal phases extracted by the narrow band filtering and application of the Hilbert transform is conceptually very different from a TE as it reduces the dimensions of state spaces of the variables entering the TE computation to two.

Another important difference of the approach proposed here and all previous approaches is the recognition of the problem of frequency resolved TE as a PID-problem—warranting particular care in the interpretation of results as laid out above.

### Resampling Methods and the free parameters

In this section we provide a description on the resampling methods implemented in the spectral mTE algorithm to shuffle the wavelet coefficients for the creation of the surrogate data.

#### Resampling Block size

The block resampling technique has been used extensively in surrogate data generation (see for example [20]). In this paper, we consider a resampling block size of 1, which can be thought of as a simple random permutation of the wavelet coefficients. The block size is an input parameter of the spectral TE and it can be set by the user (e.g. the block size is set to 32 in [20]).

#### Iterative Amplitude Adjustment Fourier Transform

The IAAFT method relies mainly on the work of [22], where a detailed description of the implementation and different applications can be found. We used the same algorithm with two fundamental changes. First, we apply the IAAFT method at one scale at time. This is motivated by the necessity to destroy putative TE information one scale at a time, keeping the contribution of the other scales intact. Second, with the IAAFT we do not apply any threshold to retain wavelet coefficients intact at a certain scale (to refer to [22] we set the threshold *p* = 0, so all coefficients are randomly shuffled and go to the iterative amplitude adjustment) since our goal was not to have a qualitative analysis between surrogate data and original data.

#### Choice of Target History Coverage

Equation 1, in the subsection 2.1.1 *Multivariate Transfer Entropy*, contains the candidate source past set *S*_<*t*_ and the candidate target past set *T*_<*t*_ which have to be defined to compute mTE. In the presented simulations, we set the maximum lag of the target to cover 1/4 of the cycle of the lowest frequency of interest (e.g. if the lowest frequency of interest was 4 Hz, we covered 1/4 of the cycle of 4 Hz). The maximum source lag was set to 3 samples lag, since the true delays were known in the simulations (1 or 2 samples lag). In case of other applications the maximum source lag should span a plausible number of samples for the system under study (e.g. a range of plausible axonal conduction delays in the case of neural data).

#### A cautionary note on frequency specific information transfer versus cross frequency coupling

Before concluding, we would like to stress that our novel algorithm should not be misunderstood as an analysis technique to estimate cross-frequency coupling (CFC, see [34] and references therein). This is because, first, information transfer and coupling are conceptually different (and it is transfer that is more important when trying to understand a computation, whereas coupling is important to understand the biophysics and dynamics of a system, [36,41]). We also note again that the concept of information being transferred from individual source frequencies to individual target frequencies—as it is expressed in the specific wording ‘cross-frequency’—does not reflect the actual complexity of the problem.

## Conclusion

We here present an algorithm that returns the frequencies at which a source sends information to a target via any (possibly nonlinear) mechanism, or at which the target receives information from a source. We discuss that a full analysis of frequency-resolved information transfer is a problem of the partial-information decomposition type, such that results should be interpreted carefully, and in the light of possible synergies and redundancies between frequencies in the source or the target. Against this background, we also present a test for a potential information transfer from a source frequency to a target frequency. While our algorithms are motivated by problems from neuroscience they are applicable in all fields where frequency-specific information transfer is of interest, e.g. turbulence or climate research.

## Acknowledgements

We are grateful to Christopher J. Keylock for introducing the original idea of combining TE with IMODWT-generated surrogate data to us, and for fuitful discussions on the topic. EP, OT, MW received support from the SFB 1193, C04, funded by the DFG. MW is a professor at the Campus Institute for Dynamics of Biological Networks (CIDBN) funded by the Volkswagen Stiftung.

## 1 Supplementary Material

### 1.1 Information transfer from one source frequency to multiple target frequencies (LA16)

**Fig S1.**
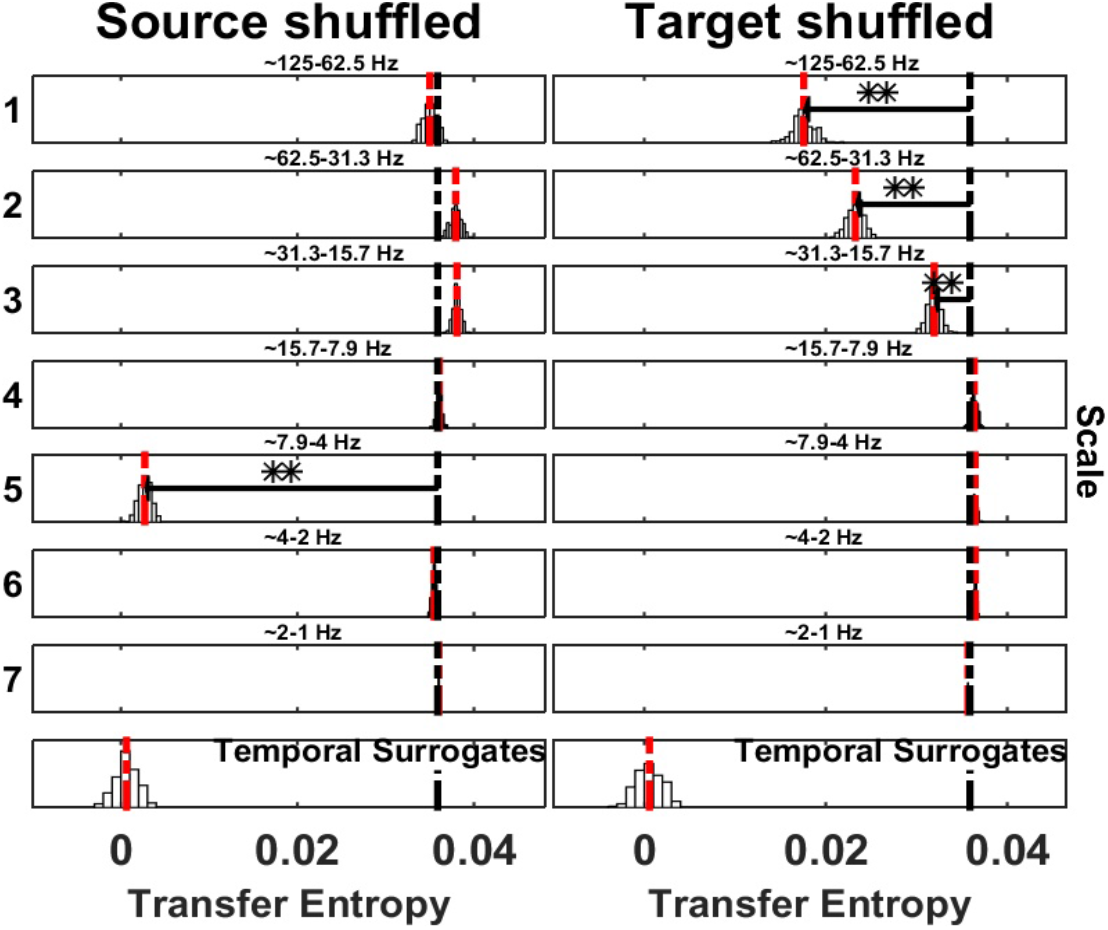
Spectrally resolved Transfer Entropy. See Fig. 5 for display conventions. (Left panel) Information transfer, drops when wavelet coefficients are selectively shuffled at scale 5 (frequency band 4-8 Hz) on the source site. The corresponding reception of information at the target is shown on the right panel, where a drop for shuffled wavelet coefficients is observed at scale 1 (frequency band 63-125 Hz), scale 2 (frequency band 31-63 Hz) and scale 3 (frequency band 16-31 Hz)

## References

1. Schreiber T. Measuring information transfer. Physical Review Letters. 2000;85(2):461–464. doi:10.1103/PhysRevLett.85.461.

2. Buzsaki G. Rhythms of the Brain. Oxford University Press; 2006.

3. Besserve M, Scholkopf B, Logothetis NK, Panzeri S. Causal relationships between frequency bands of extracellular signals in visual cortex revealed by an information theoretic analysis. J Comput Neurosci. 2010;.

4. Barnett L, Barrett AB, Seth AK. Granger causality and transfer entropy Are equivalent for gaussian variables. Physical Review Letters. 2009;103(23). doi:10.1103/PhysRevLett.103.238701.

5. Florin E, Gross J, Pfeifer J, Fink GR, Timmermann L. The effect of filtering on Granger causality based multivariate causality measures. NeuroImage. 2010;50(2):577–588. doi:10.1016/j.neuroimage.2009.12.050.

6. Weber I, Florin E, Von Papen M, Timmermann L. The influence of filtering and downsampling on the estimation of transfer entropy. PLoS ONE. 2017;12(11):1–28. doi:10.1371/journal.pone.0188210.

7. Barnett L, Seth AK. Behaviour of Granger causality under filtering: Theoretical invariance and practical application. Journal of Neuroscience Methods. 2011;201(2):404–419. doi:10.1016/j.jneumeth.2011.08.010.

8. Vicente R, Wibral M, Lindner M, Pipa G. Transfer entropy—a model-free measure of effective connectivity for the neurosciences. Journal of computational neuroscience. 2011;30(1):45–67.

9. Ragwitz M, Kantz H. Markov models from data by simple nonlinear time series predictors in delay embedding spaces. Physical Review E. 2002;65(5):056201.

10. Wibral M, Pampu N, Priesemann V, Siebenhhner F, Hannes S, Lindner M, et al. Measuring Information-Transfer Delays. PLoS ONE. 2013;8(2).

11. Novelli L, Wollstadt P, Mediano P, Wibral M, Lizier JT. Large-scale directed network inference with multivariate transfer entropy and hierarchical statistical testing. Network Neuroscience. 2019;3(3):827–847. doi:10.1162/netn_a_00092.

12. Lizier JT. The local information dynamics of distributed computation in complex systems. Springer Science & Business Media; 2012.

13. Wollstadt P, Meyer U, Wibral M. A graph algorithmic approach to separate direct from indirect neural interactions. PLoS ONE. 2015;10(10):1–26. doi:10.1371/journal.pone.0140530.

14. Welch WJ. Algorithmic Complexity: Three NP-Hard Problems in Computational Statistics. Journal of Statistical Computation and Simulation. 1982;15(1):17–25. doi:10.1080/00949658208810560.

15. Wollstadt P, Lizier J, Vicente R, Finn C, Martinez-Zarzuela M, Mediano P, et al. IDTxl: The Information Dynamics Toolkit xl: a Python package for the efficient analysis of multivariate information dynamics in networks. Journal of Open Source Software. 2019;4(34):1081. doi:10.21105/joss.01081.

16. Lizier JT, Rubinov M. Multivariate construction of effective computational networks from observational data. Max Planck Institute: Preprint. 2012;.

17. Faes L, Nollo G, Porta A. Information-based detection of nonlinear Granger causality in multivariate processes via a nonuniform embedding technique. Physical Review E - Statistical, Nonlinear, and Soft Matter Physics. 2011;83:1–15. doi:10.1103/PhysRevE.83.051112.

18. Takens F. Detecting strange attractors in turbulence. Dynamical Systems and Turbulence, Lecture Notes in Mathematics. 1981;898:366–381. doi:10.1007/bfb0091924.

19. Lancaster G, Iatsenko D, Pidde A, Ticcinelli V, Stefanovska A. Surrogate data for hypothesis testing of physical systems. Physics Reports. 2018;748:1–60. doi:10.1016/j.physrep.2018.06.001.

20. Breakspear M, Brammer M, Robinson PA. Construction of multivariate surrogate sets from nonlinear data using the wavelet transform. Physica D: Nonlinear Phenomena. 2003;182(1-2):1–22. doi:10.1016/S0167-2789(03)00136-2.

21. Keylock CJ. Constrained surrogate time series with preservation of the mean and variance structure. Physical Review E - Statistical, Nonlinear, and Soft Matter Physics. 2006;73(3):2–5. doi:10.1103/PhysRevE.73.036707.

22. Keylock CJ. Characterizing the structure of nonlinear systems using gradual wavelet reconstruction. Nonlinear Processes in Geophysics. 2010;17(6):615–632. doi:10.5194/npg-17-615-2010.

23. Percival DBa. Wavelet methods for time-series analysis. Cambridge University Press; 2013.

24. Percival DB. Analysis of geophysical time series using discrete wavelet transforms: An overview. Lecture Notes in Earth Sciences. 2008;112:61–79. doi:10.1007/978-3-540-78938-3_4.

25. Cornish CR, Bretherton CS, Percival DB. Maximal overlap wavelet statistical analysis with application to atmospheric turbulence. Boundary-Layer Meteorology. 2006;119(2):339–374. doi:10.1007/s10546-005-9011-y.

26. Zhang Z, Telesford QK, Giusti C, Lim KO, Bassett DS. Choosing wavelet methods, filters, and lengths for functional brain network construction. PLoS ONE. 2016;11(6). doi:10.1371/journal.pone.0157243.

27. Dupré la Tour T, Tallot L, Grabot L, Doyère V, van Wassenhove V, Grenier Y, et al. Non-linear auto-regressive models for cross-frequency coupling in neural time series. PLoS Computational Biology. 2017;13(12):1–32. doi:10.1371/journal.pcbi.1005893.

28. Hyafil A. Disharmony in neural oscillations. Journal of Neurophysiology. 2017;118(1):1–3. doi:10.1152/jn.00026.2017.

29. Vakorin VA, Krakovska OA, McIntosh AR. Confounding effects of indirect connections on causality estimation. Journal of Neuroscience Methods. 2009;184(1):152–160. doi:10.1016/j.jneumeth.2009.07.014.

30. Brodski-Guerniero A, Paasch GF, Wollstadt P, Ozdemir I, Lizier JT, Wibral M. Information-Theoretic Evidence for Predictive Coding in the Face-Processing System. The Journal of neuroscience: the official journal of the Society for Neuroscience. 2017;37(34):8273–8283. doi:10.1523/JNEUROSCI.0614-17.2017.

31. Wollstadt P, Sellers KK, Rudelt L, Priesemann V, Hutt A, Fröhlich F, et al. Breakdown of local information processing may underlie isoflurane anesthesia effects. PLoS Computational Biology. 2017;13(6):1–35. doi:10.1371/journal.pcbi.1005511.

32. Maris E, Oostenveld R. Nonparametric statistical testing of EEG- and MEG-data. Journal of Neuroscience Methods. 2007;164(1):177–190. doi:10.1016/j.jneumeth.2007.03.024.

33. Bastos AM, Vezoli J, Bosman CA, Schoffelen JM, Oostenveld R, Dowdall JR, et al. Visual areas exert feedforward and feedback influences through distinct frequency channels. Neuron. 2015;85(2):390–401. doi:10.1016/j.neuron.2014.12.018.

34. Aru J, Aru J, Priesemann V, Wibral M, Lana L, Pipa G, et al. Untangling cross-frequency coupling in neuroscience. Current Opinion in Neurobiology. 2015;31(September 2014):51–61. doi:10.1016/j.conb.2014.08.002.

35. Lizier J, Bertschinger N, Jost J, Wibral M. Information decomposition of target effects from multi-source interactions: perspectives on previous, current and future work; 2018.

36. Wibral LJT Michael, Vicente R. Directed Information Measures in Neuroscience. Springer; 2014.

37. Paluš M. Coupling in complex systems as information transfer across time scales. Philosophical Transactions of the Royal Society A: Mathematical, Physical and Engineering Sciences. 2019;377. doi:http://doi.org/10.1098/rsta.2019.0094.

38. Shi W, Yeh C, Hong Y. Cross-Frequency Transfer Entropy Characterize Coupling of Interacting Nonlinear Oscillators in Complex Systems. IEEE Transactions on Biomedical Engineering. 2019;66(2):521–529. doi:10.1109/TBME.2018.2849823.

39. Lobier M, Siebenhühner F, Palva S, Palva JM. Phase transfer entropy: A novel phase-based measure for directed connectivity in networks coupled by oscillatory interactions. NeuroImage. 2014;85:853–872. doi:10.1016/j.neuroimage.2013.08.056.

40. Xu S, Baldea M, Edgar TF, Wojsznis W, Blevins T, Nixon M. Root cause diagnosis of plant-wide oscillations based on information transfer in the frequency domain. Industrial & Engineering Chemistry Research. 2016;55(6):1623–1629.

41. Lizier JT, Prokopenko M, Zomaya AY. Local measures of information storage in complex distributed computation. Information Sciences. 2012;208:39–54. doi:10.1016/j.ins.2012.04.016.

