## Supplementary Figure S1 for "Measuring spectrally-resolved information transfer for sender- and receiver-specific frequencies"

### 1 Supplementary Material

#### 1.1 Information transfer from one source frequency to multiple target frequencies (LA16)

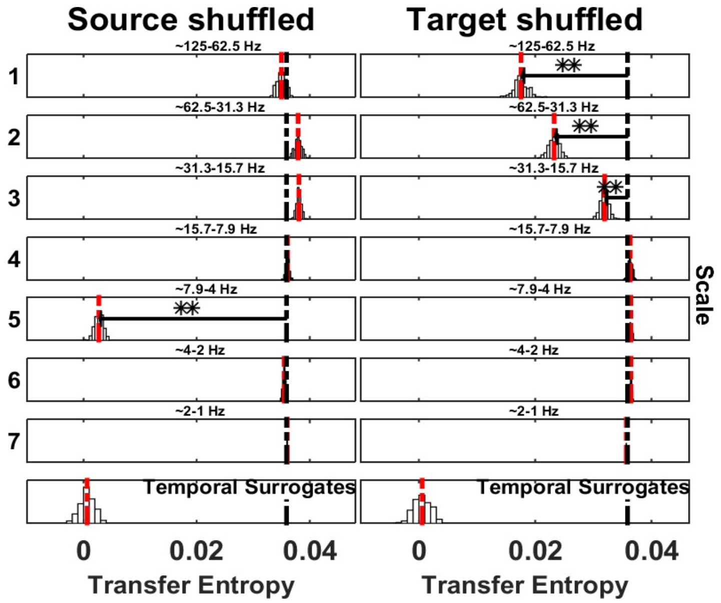


**Fig S1. Spectrally resolved Transfer Entropy**. See Fig. 5 for display conventions. (Left panel) Information transfer, drops when wavelet coefficients are selectively shuffled at scale 5 (frequency band 4-8 Hz) on the source site. The corresponding reception of information at the target is shown on the right panel, where a drop for shuffled wavelet coefficients is observed at scale 1 (frequency band 63-125 Hz), scale 2 (frequency band 31-63 Hz) and scale 3 (frequency band 16-31 Hz)
